# Temporal regulation of Pten is essential for retina regeneration in zebrafish

**DOI:** 10.1101/2021.07.09.451767

**Authors:** Shivangi Gupta, Poonam Sharma, Mansi Chaudhary, Sharanya Premraj, Simran Kaur, V Vijithkumar, Rajesh Ramachandran

## Abstract

Unlike mammals, zebrafish possess a remarkable ability to regenerate damaged retina after an acute injury. Retina regeneration in zebrafish involves the induction of Müller glia-derived progenitor cells (MGPCs) exhibiting stem cell-like characteristics, which are capable of restoring all retinal cell-types. Here, we explored the importance of Phosphatase and tensin homolog (Pten), a dual-specificity phosphatase and tumor suppressor during retina regeneration. The Pten undergo rapid downregulation in the Müller glia and is absent in MGPCs, which is essential to trigger Akt-mediated cell proliferation to cause retina regeneration. We found that the forced downregulation of Pten accelerates MGPCs formation, while its overexpression restricts the regenerative response. We observed that Pten regulates the proliferation of MGPCs not only through Akt pathway but also by Mmp9/Notch signaling. Mmp9-activity is essential to induce the proliferation of MGPCs in the absence of Pten. Lastly, we show that Pten expression is fine-tuned through Mycb/histone deacetylase1 and Tgf-β signaling. The present study emphasizes on the stringent regulation of Pten and its crucial involvement during the zebrafish retina regeneration.

## Introduction

Retina regeneration is an enigmatic phenomenon in which the functional restoration of damaged parts occurs in vertebrates such as zebrafish, unlike found in the mammals. Several studies were carried out using the zebrafish model that deciphered the significance and necessity of various gene regulatory events during retina regeneration (Goldman, 2014; Lahne et al., 2020; Wan and Goldman, 2016). An acute injury to zebrafish retina induces the Müller glia (MG) to generate Müller glia-derived retinal progenitor cells (MGPCs), capable of giving rise to all retinal cell types (Hamon et al., 2016; Lahne et al., 2020). The study of regenerative response of retina in mammals, birds, and fishes have pointed out important similarities and differences among these model organisms (Gallina et al., 2014). Further, the knowledge from zebrafish retina regeneration studies is far from complete to take it to the mammalian models, and eventually humans.

Several past studies on tissue regeneration have focused on the importance of transcription factors, chromatin/epigenome modifiers, pluripotency inducing factors, and various cellular signaling events (Goldman, 2014; Lahne et al., 2020; Wan and Goldman, 2016). Albeit with knowledge on the involvement of calcineurin, a calcium-dependent serine-threonine phosphatase, during fin regeneration in zebrafish (Kujawski et al., 2014; McMillan et al., 2018), little is known about the importance of dual-specificity phosphatases during retina regeneration. Unlike kinases, the protein phosphatases have a broad range of substrates, making them a less attractive candidate for exploration. Nonetheless, the knowledge of phosphatases during tissue regeneration is pivotal in harnessing the proliferative responses of the progenitors. Phosphatases have roles in regulating various molecular pathways such as STAT3 signaling (Kim et al., 2018), VEGF signaling (Corti and Simons, 2017), Ras/ERK signaling (Kidger and Keyse, 2016), and also in cancer (Ostman et al., 2006; Pestell et al., 2000; Pulido and Hooft van Huijsduijnen, 2008), which make them attractive candidates to unravel mysteries of regeneration biology.

To fill in the missing parts of the jigsaw puzzle of regeneration, we explored the significance of a unique tumor suppressor gene, phosphatase and tensin homolog (Pten), which is also a dual-specificity phosphatase (Dahia, 2000). PTEN is a versatile molecule implicated in various biological phenomena such as tumor suppression (Li and Ross, 2007; Simpson and Parsons, 2001), DNA damage repair (Ming and He, 2012), neurological disorders such as autism (Rademacher and Eickholt, 2019), cell-size regulation (Backman et al., 2002), and regulation of chromatin dynamics (Yang and Yin, 2020). PTEN, as a dual-specificity phosphatase (Dahia, 2000; Eng, 1999), acts both in cytoplasm and nucleus (Planchon et al., 2008). PTEN is important to maintain the retinal architecture and function (Cantrup et al., 2012; Jo et al., 2012; Kim et al., 2008; Sakagami et al., 2012; Tachibana et al., 2016). Pharmacological inhibition of PTEN has been proposed as a strategy to enhance tissue regeneration (Borges et al., 2020). Despite the knowledge on the relevance of PTEN during axon regeneration (Kurimoto et al., 2010; Mak et al., 2020; Ohtake et al., 2015), its importance during retina regeneration remained underexplored.

In this study of retina regeneration, we explored the significance of Pten regulation and its downstream molecular events, which are well-elucidated in various tumorigenic pathways. We find rapid regulation of Pten soon after retinal injury. Therefore, we investigated further the mechanistic involvement of Pten during zebrafish retina regeneration. Subsequently, we hypothesized that the MG dedifferentiation could depend on Pten-mediated signaling and may have some similarities to the gene expression events during cancer progression. Since we were interested in the possible involvement of Pten during the immediate early response of MG to injury, we analyzed the retina upon blockade of Pten. We identified that the expression pattern of several essential genes associated with retina regeneration was induced by Pten-mediated cellular signaling and *vice versa* that reveal the stringent gene regulatory network associated with retina regeneration. These include the interplay of the Pten/PI3K/Akt pathway, Pten/Notch/Tgf-β signaling components, and matrix metalloproteinase 9 (Mmp9). The findings from this study shed light on the existing knowledge of regeneration by unraveling the Pten-mediated gene regulatory events and its regulation through a plethora of other factors.

## Results

### Pten is induced soon after injury, but not in the MGPCs

We explored the expression pattern of two genes of Pten, namely *ptena* and *ptenb* in zebrafish retina after an acute injury. The Ptena and Ptenb share 80% similarity in their amino acid sequences (Figure EV1A), and are known to play distinct roles during zebrafish embryogenesis (Croushore et al., 2005). The mRNAs of *ptena* and *ptenb* were quantified at various times post retinal injury. We found a dual peak of induction of *ptena* (Figure 1A), and *ptenb* (Figure 1B) with a larger induction soon after injury which marks the dedifferentiation phase, followed by a short peak during the proliferative phase around 4dpi compared to the uninjured control. We then assessed the qualitative expression of *ptena* and *ptenb* in the injured retina at different time points after retinal injury, such as 2, 4 and 6dpi (Figures 1C, 1E, EV1B, and EV1C). The expression of *pten* genes seemed to be secluded from the proliferating population of cells, which are deemed to be the MG derived progenitor cells (MGPCs) as revealed by mRNA *in situ* hybridization performed in injured retinal sections. Quantitative analysis revealed a significantly reduced co-labeling of *pten* mRNAs with proliferating cell nuclear antigen (PCNA) marker at 2, 4, and 6dpi (Figures 1D and 1F). Interestingly, the cells flanking the PCNA^+^ MGPCs seemed to express more *pten* mRNAs. This observation supported the already reported anti-proliferative signaling events emanated from Pten in stem cells and cancer (Hill and Wu, 2009; Luongo et al., 2019; Rossi and Weissman, 2006; Stumpf et al., 2015). We then explored if the observed exclusion of *ptena*/*ptenb* mRNA from the MGPCs was reflected in their protein levels as well. While the Pten levels were fairly uniform in expression pattern in uninjured retinal cells, we saw a drastic decline in its levels in the MGPCs at 4dpi (Figure 1G and EV1D). To affirm the expression of Pten in the MG cells, we also used *gfap*:GFP transgenic retina, which marks the MG cells with GFP. We saw a co-expression of Pten and GFP in the uninjured *gfap*:GFP transgenic retina (Figure EV1E). A significant number of PCNA^+^ MGPCs had low levels of Pten (Figure 1H). Further to validate this, we took advantage of a transgenic zebrafish line *1016tuba1a*:GFP, which could label the MGPCs with GFP after an acute injury (Fausett and Goldman, 2006; Kaur et al., 2018; Mitra et al., 2018; Mitra et al., 2019; Sharma et al., 2019; Sharma et al., 2020) and explored the expression pattern of Pten. We found a significant decline in the Pten expression in GFP^+^ cells in the retina at 4dpi, both qualitatively (Figure 1I), and quantitatively (Figure 1J). It was further validated by qPCR analysis of *pten* mRNA levels in sorted cells from *1016tuba1a*:GFP transgenic retina (Figures EV1F and EV1G). Despite the reduced Pten levels in the MGPCs, there was no appreciable change in the Pten levels in the total retina (Figures 1K and 1L), probably because of reduced representation of MGPCs formed in the retina through needle stab injury (Ramachandran et al., 2012). These observations support the anti-proliferative role of Pten as a tumor suppressor protein as reported in other systems.

**Figure 1:**
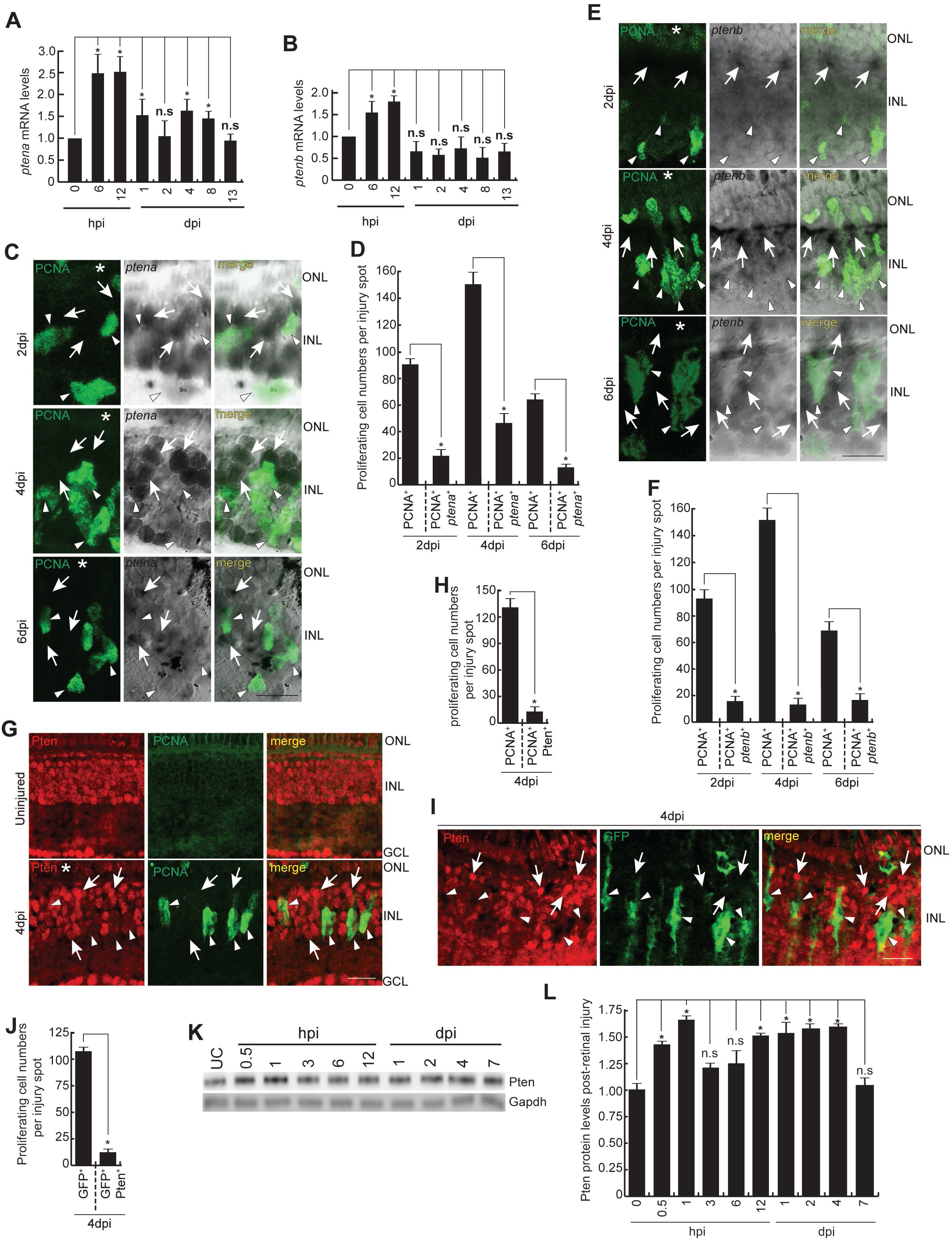
Pten is expressed in the whole retina and is absent in the MGPCs. (**A** and **B**) The qPCR analyses of *ptena* (**A**) and *ptenb* (**B**) genes in the retina at various time points post-retinal injury; *p < 0.04; n=6 biological replicates. (**C** and **D**) The *in-situ* hybridisation (ISH) and Immunofluorescence (IF) microscopy images of retinal cross-sections show the expression of *ptena* mRNA in the neighbouring cells of PCNA^+^ MGPCs at 2dpi, 4dpi and 6dpi (**C**), which is quantified (**D**); *p < 0.0001; n=6 biological replicates. (**E** and **F**) ISH and IF microscopy images of retinal cross-sections reveal the expression of *ptenb* mRNA in the neighbouring cells of PCNA^+^ MGPCs at 2dpi, 4dpi and 6dpi (**E**), which is quantified (**F**); *p < 0.0001; n=6 biological replicates. (**G** and **H**) IF microscopy images of retinal cross-sections show the pan-retinal expression of Pten in the uninjured retina, while being almost absent in the PCNA^+^ MGPCs in 4dpi retina (**G**), which is quantified (**H**); *p < 0.00005; n=6 biological replicates. (**I** and **J**) IF microscopy images of a retinal cross-section show highly reduced expression of Pten in the GFP^+^ MGPCs of *1016tuba1a*:GFP transgenic fish retina at 4dpi (**I**), which is quantified (**J**); *p < 0.005; n=6 biological replicates. (**K** and **L**) Western Blot analysis (**K**) of Pten from retinal lysates prepared at different time points post-injury, which is quantified by densitometry (**L**); *p < 0.001; n=6 biological replicates. Gapdh is the loading control. Scale bars represent 10μm in (**C, E, G, I**); the asterisk marks the injury site and GCL, ganglion cell layer; INL, inner nuclear layer; ONL, outer nuclear layer (**C, E, G, I**); hpi, hours post injury; dpi, days post injury; white arrowheads mark PCNA^+^/*ptena*^−^ (**C**), PCNA^+^/*ptenb*^−^ (**E**), PCNA^+^/Pten^−^ (**G**), GFP^+^/Pten^−^ (**I**) cells; white arrows mark *ptena*^+^/PCNA^−^ (**C**), *ptenb*^+^/PCNA^−^ (**E**), Pten^+^/PCNA^−^ (**G**) and Pten^+^/GFP^−^ (**I**) cells; UC-Uninjured control (**K**); Error bars represent SD; n.s., not significant.

### Downregulation of Pten increases the number of MGPCs during retina regeneration

We then explored the effects of downregulation of Pten during retina regeneration. For this, we employed a morpholino (MO)-based gene knockdown approach, an efficient and widely used strategy in zebrafish retina (Mitra et al., 2018; Mitra et al., 2019; Sharma et al., 2019; Sharma et al., 2020). MOs against *ptena* and *ptenb* were electroporated in retina at different concentrations at the time of injury and probed for the proliferation of MGPCs at 4dpi (Figure 2A). The random delivery of lissamine fluorescent-tagged MOs into the retinal cells blocks the production of Pten protein. With the knockdown of *ptena* and *ptenb*, we found a MO dose-dependent increase in the number of BrdU-labelled MGPCs in 4dpi retina compared to standard control MO or *ptena*/*ptenb* mismatch MOs (Figures 2B and 2C). Double knockdown of *ptena* and *ptenb* had a more profound effect on the number of MGPCs (Figure EV2A and EV2B). We also performed rescue experiments for *ptena* MO and *ptenb* MO to rule out their off-target effects, using their corresponding mRNAs (Figures EV2C-EV2F). Overexpression of *ptena* and *ptenb* mRNA resulted in a drastic decline in the number of MGPCs in 4dpi retina (Figures EV2G and EV2H).

**Figure 2:**
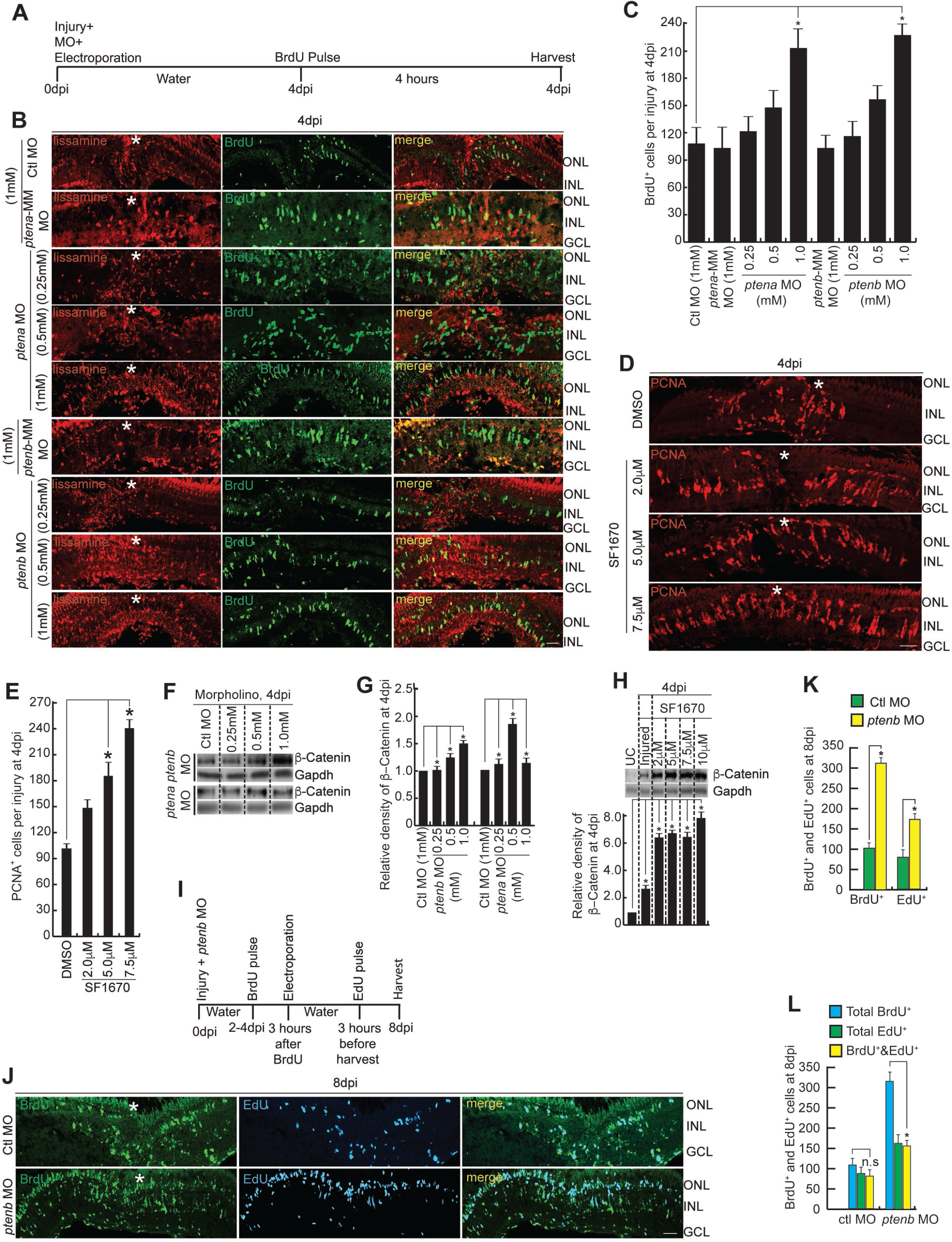
Downregulation and blockade of Pten enhances proliferation of MGPCs. (**A**) An experimental timeline that describes injury, Morpholino (MO) delivery, electroporation at 0dpi, BrdU pulse for 4hrs at 4dpi, followed by harvesting. (**B** and **C**) IF microscopy images of retinal cross-sections show an increase in the number of BrdU^+^ MGPCs with the increasing concentrations of *ptena* and *ptenb* MOs (Lissamine tag), compared to control MO/*ptena*/*ptenb*-mismatch (MM) MO-injected retina at 4dpi (**B**), which is quantified (**C**); *p < 0.04; n=9 biological replicates. (**D** and **E**) IF microscopy images of retinal cross-sections show an increase in the number of PCNA^+^ MGPCs with the increasing concentrations of SF1670 at 4dpi (**D**), which is quantified (**E**); *p < 0.03; n=6 biological replicates. (**F** and **G**) Western Blot analyses of β-catenin in retinal extracts collected after *ptena* and *ptenb* knockdown in retinae at 4dpi (**F**), quantified by densitometry (**G**); *p < 0.001; n=6 biological replicates. (**H**) Western Blot analysis of β-catenin in retinal extracts prepared from retinae treated with different concentrations of SF1670 at 4dpi (above), quantified by densitometry (below); *p < 0.001; n=6 biological replicates. (**I**) An experimental timeline that describes *ptenb* MO injection at the time of injury, BrdU pulse-labeling for 4hrs at 2-4dpi followed by electroporation on 4dpi done 3hrs post BrdU exposure, and harvesting at 8dpi after 3hrs of EdU pulse. (**J-L**) IF microscopy images of retinal cross-sections show an increase in the number of BrdU^+^ and EdU^+^ MGPCs in *ptenb* MO-injected retina as compared to the Control MO-injected retina at 8dpi (**J**), which is quantified (**K** and **L**); *p < 0.04; n=4 biological replicates. Scale bars represent 10μm in (**B, D, J**); the asterisk marks the injury site and GCL, ganglion cell layer; INL, inner nuclear layer; ONL, outer nuclear layer (**B, D, J**); dpi, days post injury; Error bars represent SD; Gapdh is the loading control (**F, H**) UC-Uninjured control (**H**); Ctl MO-Control morpholino (**B,C,F,G,J,K**); n.s., not significant.

Similar results were observed when Pten activity was blocked using a pharmacological inhibitor SF1670 (Spinelli et al., 2015; Wang et al., 2020) in regenerating retina at 4dpi (Figures 2D and 2E). The *1016tuba1a*:GFP transgenic zebrafish retina also had an elevated GFP expression because of SF1670 treatment (Figure EV2I), which supports the enhanced the proliferation of MGPCs seen in the wild-type retina. It is also interesting to note that the uninjured retina did not show any proliferative response to SF1670 treatment (Figure EV2J). Furthermore, the Pten blockade by dipping the fish in SF1670 during the dedifferentiation and proliferation phases, 0-2dpi and 2-4dpi respectively, caused an enhanced number of MGPCs (Figures EV3A and EV3B). This result suggests that the inhibition of Pten caused an enhanced number of MGPCs irrespective of the regeneration phase. The blockade of Pten did not lead to apoptosis, as revealed in the TUNEL assay (Figure EV3C). The increase in MGPCs because of *pten* knockdown is also reflected in the phospho-histone positive cells, indicative of active mitosis (Figures EV2H-now Figure EV3D and EV3E).

PTEN is known to interfere with the stability of glycogen synthase kinase 3 beta (GSK3β) (Huang et al., 2007; Korkaya et al., 2009), an enzyme known for its destabilization effect on pro-proliferative molecule β-catenin, through Phosphatidylinositol 3’-kinase (PI3K)/Akt axis in several conditions including cancer (Lu et al., 1999). Increased β-catenin levels have been associated with the proliferation of MGPCs in regenerating zebrafish retina (Ramachandran et al., 2011). We explored if the downregulation of Pten affected the β-catenin protein levels, which may account for the increased number of MGPCs. We saw a *ptena/ptenb* MO or SF1670 concentration-dependent increase in the β-catenin levels (Figures 2F, 2G, 2H, EV3F, and EV3G), suggestive of the functional existence of a Pten-PI3K/Akt/β-catenin signaling axis during zebrafish retina regeneration. Further, we traced the lineage of the enhanced MGPCs in Pten blockade with SF1670. These increased MGPCs found in SF1670-treated retina were able to give rise to major retinal cell-types when analyzed at 30dpi indicative of their ability to differentiate to various cell types (Figures EV3H and EV3I).

We probed further to find if the observed increase in MGPCs because of Pten inhibition is due to more MGs entering cell cycle or the MGPCs failing to exit the cell cycle. To address this, the retina was allowed to regenerate until 4dpi, a time of peak proliferation of MGPCs, and then the Pten activity was blocked during 4-8dpi, a time when the proliferation reduces. During this descent phase, Pten was blocked to address if a similar effect was seen on the proliferation of MGPCs as it was seen with continuous Pten blockade during 0-4dpi. We adopted a double-labeling approach (Figure 2I), in which most of the MGPCs were labeled with BrdU from 2-4dpi followed by Pten inhibition, and a final EdU labeling before harvest at 8dpi. In this approach, we could differentially label the MGPCs from 4dpi and the newly formed ones, if any, at 8dpi. If the BrdU labeled MGPCs continue to divide until 8dpi, they would contain both BrdU and EdU, and if the observed increase in MGPCs is because of new progenitors, then the cells should contain only EdU. Interestingly, we saw a significantly co-labeled BrdU^+^ and EdU^+^ cells at 8dpi, suggestive of the first option where the right proportion of MGPCs continued to be in cell-cycle in the absence of Pten causing an overall increase in the number of BrdU^+^ and EdU^+^ cells (Figures 2J-2L). However, it is important to note that in *ptenb* MO-electroporated conditions, the total number of BrdU cells were higher than EdU and BrdU/EdU co-labeled cells (Figure 2L). This observation suggests that upon *ptenb*-late knockdown, cells continued to proliferate for some more time and later started exiting the cell cycle owing to a reduced number of BrdU/EdU positive cells as compared to the total number of BrdU^+^ cells. These results support the idea that the presence of Pten in the neighboring cells of MGPCs is essential to curb the hyper-proliferative response during retina regeneration.

### Downregulation of Pten activates Akt

Despite being a protein phosphatase, the actions of Pten mainly occur when it is membrane-bound, wherein it dephosphorylates the phosphatidylinositol triphosphate (PIP3) to PIP2 (Das et al., 2003). The reverse reaction is done by another enzyme PI3K. Pten is also known to interfere with PI3K activity causing a double impact on the membrane-bound PIP3 (Carracedo and Pandolfi, 2008). The membrane-bound PIP3 recruits two other enzymes, namely phosphoinositide-dependent kinase 1 (PDK1) and protein kinase B (PKB, also known as Akt), causing the phosphorylation and activation of the Akt by PDK1 (Vanhaesebroeck and Alessi, 2000). This Pten-PI3K-Akt axis is implicated in various biological processes such as cell proliferation, DNA repair, apoptosis (Yin et al., 2014), and diseases such as cancer (Carnero et al., 2008; Nathan et al., 2017). Since Pten was absent in the MGPCs, we anticipated an upregulation of Akt in the proliferating cells. We explored the expression pattern of Akt in regenerating zebrafish retina and found a co-labeling of Akt in the PCNA^+^ MGPCs in 4dpi retina, and no appreciable expression in the uninjured conditions (Figure 3A). The Akt also seemed to be in its biologically active form, being phosphorylated at both Serine467 (Ser473 in mammals) and Threonine302 (Thr308 in mammals) amino acids (Cheng et al., 2013), in PCNA^+^ MGPCs in 4dpi retina (Figures 3B and 3C). We then explored if the trend remains the same with forced inhibition of Pten activity either through SF1670 or *ptena*/*ptenb* MOs. We saw an increase in the Akt and its phosphorylated forms in 4dpi retinal samples treated continuously with SF1670 (Figures 3D and 3E). Similar trends were also seen in the levels of Akt and its phosphorylated forms with Pten blockade in 16hpi and 2dpi retinal extracts (Figures EV4A-EV4D). Moreover, upon panretinal NMDA-mediated injury to the retina, a strong proliferative response in the entire retina was observed (Figure EV4E), unlike upon focal mechanical injury. Interestingly, it showed similar trends of exclusion of Pten expression from the MGPCs (Figure EV4F) and the increased expression of Akt and its phosphorylated forms (Figures EV4G and EV4H), as was seen upon mechanical retinal injury at 4dpi. These observations support the view that the gene expression paradigm in a regenerating zebrafish retina is conservative irrespective of the mode of injury as previously reported (Powell et al., 2016).

**Figure 3:**
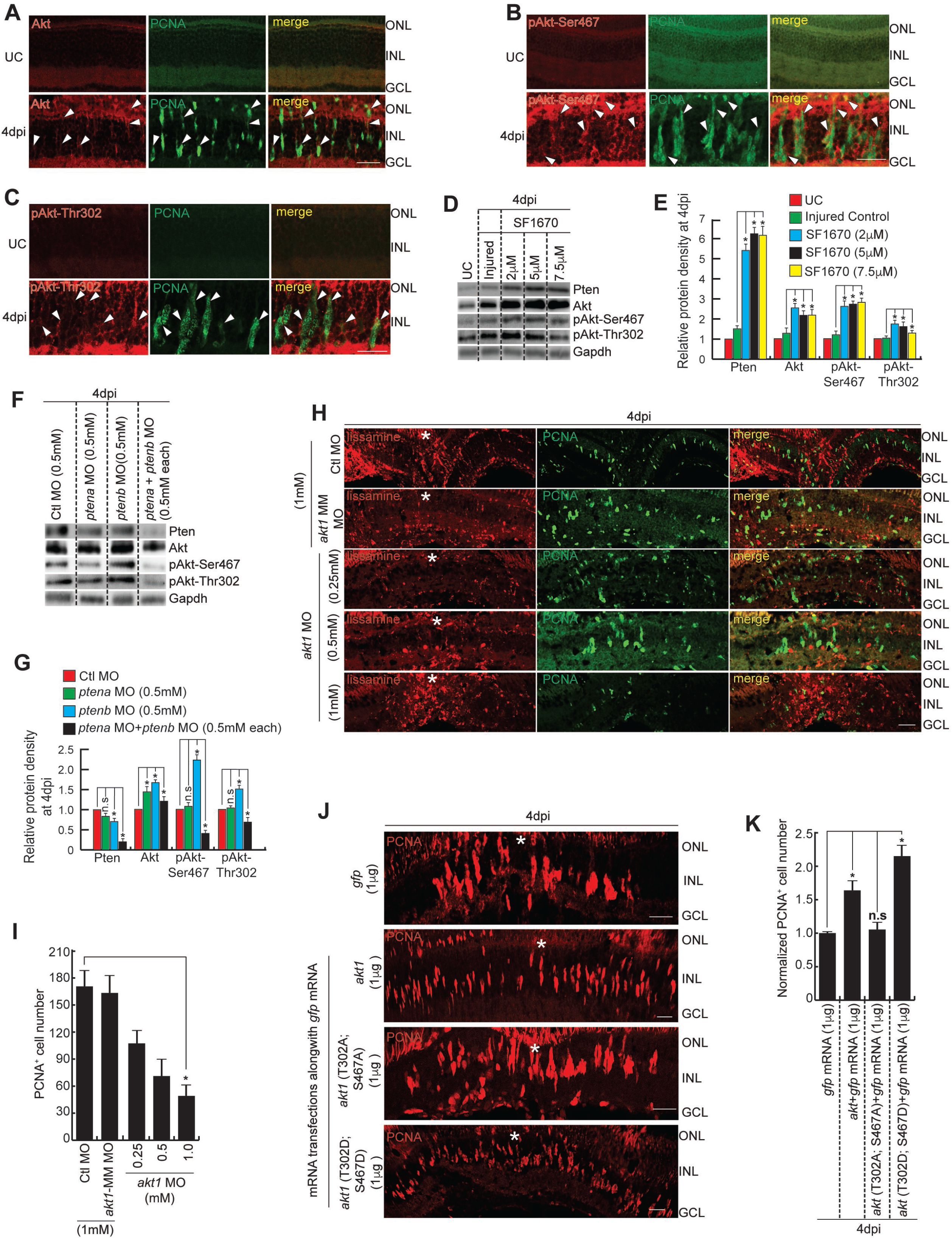
Downregulation of Pten activates Akt which increases the number of MGPCs. (**A-C**) IF microscopy images of retinal cross-sections show the expression of Akt (**A**), pAkt-Ser467 (**B**) and pAkt-Thr302 (**C**) in the PCNA^+^ MGPCs in 4dpi retina, while being absent in uninjured retina. (**D** and **E**) Western Blot analyses of Pten, Akt, pAkt-Ser467, pAkt-Thr302 from retinal extracts prepared from retinae injected with different concentrations of SF1670 at 4dpi (**D**), quantified by densitometry (**E**); *p < 0.001; n=6 biological replicates. (**F** and **G**) Western Blot analyses of Pten, Akt, pAkt-Ser467, pAkt-Thr302 from retinal extracts collected after *ptena*/ *ptenb* knockdown in retinae at 4dpi (**F**), quantified by densitometry (**G**); *p < 0.001; n=6 biological replicates. (**H** and **I**) IF microscopy images of retinal cross-sections show a decline in the number of PCNA^+^ MGPCs with the increasing concentrations of *akt1* MO compared to control MO/*akt1* MM MO-injected retina at 4dpi (**H**), which is quantified (**I**); *p < 0.04; n=6 biological replicates. (**J** and **K**) IF microscopy images of retinal cross-sections show the number of PCNA^+^ MGPCs upon transfection of retina with *akt1* wild-type mRNA or with phosphomimetic form of *akt1* mRNA which increases, while the cell number remains unchanged upon retinal transfection with neutral mutation-bearing *akt1* mRNA at 4dpi (**J**), as compared to *gfp*-transfected control, which is quantified (**K**); *p < 0.03; n=6 biological replicates. Scale bars represent 10μm in (**A, B, C, H, J**); the asterisk marks the injury site in (**H, J**); GCL, ganglion cell layer; INL, inner nuclear layer; ONL, outer nuclear layer (**A, B, C, H, J**); dpi, days post injury; white arrowheads mark Akt^+^/PCNA^+^ (**A**), pAkt-Ser467^+^/PCNA^+^ (**B**) and pAkt-Thr302^+^/PCNA^+^ (**C**) cells; Error bars represent SD; Gapdh is the loading control (**D, F**); UC-Uninjured control (**A-E,**); Ctl MO-Control morpholino (**F-I**); n.s., not significant.

Notably, the Pten protein itself was going up, which is biologically inactive in SF1670-treated retina, probably because of the absence of negative feedback regulation. Similar to SF1670 treatment, we observed an increase in Akt and its phosphorylated forms in *ptena*/*ptenb* MO-treated retina (Figures 3F and 3G). However, when both *ptena* and *ptenb* were knockdown simultaneously, or upon strong Pten blockade we saw a decline in the levels of phosphorylated forms of Akt. Akt is known to activate mTORC1 either indirectly via TSC1/2 inhibition or directly through silencing of mTORC1 inhibitor subunit PRAEV40 (Inoki et al., 2002; Laplante and Sabatini, 2012; Potter et al., 2002; Vander Haar et al., 2007). There exists a negative feedback regulation mechanism of mTORC1 or its downstream effectors on PI3K and mTORC2 during mice axonal regeneration (Miao et al., 2016) and in cancerous conditions (Efeyan and Sabatini, 2010). We speculated that this negative feedback regulation ensuing from mTORC1 on Akt1 phosphorylation might be the reason for the unanticipated decrease in the levels of phosphorylated Akt1 that was seen upon strong Pten blockade or combined *pten* knockdown. We observed that Rapamycin-mediated mTORC1 blockade (He et al., 2017; Zhang et al., 2020) led to a decrease in the number of MGPCs (Figures EV4I and EV4J), and increase in the levels of phosphorylation of Akt at Thr302 and Ser467 in the retina at 4dpi (Figures EV4K and EV4L). This feedback regulation may exist to maintain tissue homeostasis by preventing excess proliferation of the MGPCs due to mTORC1 activity upon Akt activation, which otherwise will get continuous ‘ON’ signals itself from activated Akt, in case of absence of this negative feedback mechanism.

Further, we explored if Akt was necessary for the normal regenerative response in the retina. For this, we blocked *akt1* mRNA using MO soon after the injury and assayed the proliferative response of MGPCs in the retina at 4dpi (Figure 3H). We saw a significant decline in the number of MGPCs compared to control MO or *akt1* mismatch MO-treated retina at 4dpi, suggestive of the necessity of Akt for retina regeneration (Figure 3I). We also performed rescue experiments for *akt1* MO to rule out its off-target effects, using its corresponding mRNA (Figures EV5A and EV5B). We then adopted an overexpression strategy of Akt by its mRNA transfection into the injured retina. For this, we used three different types of *akt1* mRNAs, namely the wild type, *akt1* mRNA with a neutral mutation to alanine of Thr302 and Ser467, and another one with phosphomimetic mutations where both Thr302 and Ser467 were mutated to aspartic acid to mimic a constitutively active form of Akt (Hart and Vogt, 2011). We saw an increase in the number of MGPCs in retina transfected with mRNA of wild-type *akt1* and its phosphomimetic form, as compared to *gfp* control mRNA and *akt1* mRNA with neutral mutation, at 4dpi (Figure 3J). Quantitative analysis showed that overexpression of Akt in native or phosphomimetic form had a positive effect on the retinal proliferation of MGPCs (Figure 3K). There were anticipated changes in the Akt and its phosphorylated forms depending on the quantity and nature of mRNA transfected (Figure EV5C and EV5D). These results emphasized the significance of the Pten-Akt regulatory axis as a major contributor to the proliferation of MGPCs during the retina regeneration.

### The involvement of PI3K and mTORC2 in retina regeneration

The Akt protein gets phosphorylated at various amino acid residues, of which Thr308 (Thr302 in zebrafish), and Ser473 (Ser467 in zebrafish) amino acids are the most important ones for its complete activation which are catalyzed by PDK1 and mTOR-complex 2 (mTORC2), respectively (Liu et al., 2014). We decided to explore whether Akt phosphorylation is necessary for retina regeneration. First, we decided to block PI3K, the activator of PDK1, to block phosphorylation at Thr308 of Akt using a drug LY294002 (Salh et al., 1998). Continuous exposure to LY294002 drug significantly abolished the PCNA^+^ MGPCs when observed at 4dpi (Figures 4A and 4B). The effect of PI3K inhibition on Akt phosphorylation was also observed in retina at 16hpi (Figures 4C and EV5E), 2dpi (Figures 4D and EV5F), and 4dpi (Figures 4E and EV5G). Similar to this, we used another drug Torin1 to block mTORC2 (Mohankumar et al., 2011) that is important in phosphorylation at Ser473 of Akt. Similar to that found with PI3K inhibition, we saw a dose-dependent reduction in the proliferating population of MGPCs in Torin1-treated retina at 4dpi (Figures 4F and 4G). We also found a reduction in the levels of phosphorylated Akt proteins at 16hpi (Figures 4H and EV5H), 2dpi (Figure 4I and EV5I), and 4dpi (Figure 4J and EV5J). Treatment of retina with LY294002 or Torin1 did not interfere with the standard apoptosis rate, as observed by the TUNEL assay (Figure EV5K). These results suggested that Akt phosphorylation is an essential step in its efficacy to mediate cellular proliferation in regenerating retina. However, we were curious to find whether the sole purpose of Pten reduction in proliferating MGPCs of regenerating retina is to activate Akt. In order to explore this, we decided to perform a double blockade experiment in which Pten was blocked using SF1670 in combination with LY294002 or Torin1 in separate experiments (Figure EV5L). The purpose of this experiment was to block Pten as well as activation of Akt and thus to block this pathway. In such a double-blocked scenario, any effect on the proliferation of MGPCs in regenerating retina would be through an Akt-independent pathway. Interestingly, in double blocker experiments, where retina was treated with SF1670 and Torin1, we saw the number of MGPCs lesser than Pten-blocked condition alone, and more than that seen in mTORC2-blocked retina and insignificant compared to DMSO control (Figures 4K and 4L). In a similar experiment where both Pten and PI3K were inhibited, we saw a persistent increase in the number of MGPCs as found with Pten inhibition by SF1670 treatment, which seemingly nullified the anti-proliferative effect observed because of PI3K inhibition by LY294002 (Figure 4M and 4N). This observation suggests that Pten-inhibition could influence the proliferation of MGPCs even in the absence of PI3K activity. These findings suggested that Pten could impact the proliferation of MGPCs through pathways other than mediated by Akt/PI3K/mTORC2 axis during retina regeneration.

**Figure 4:**
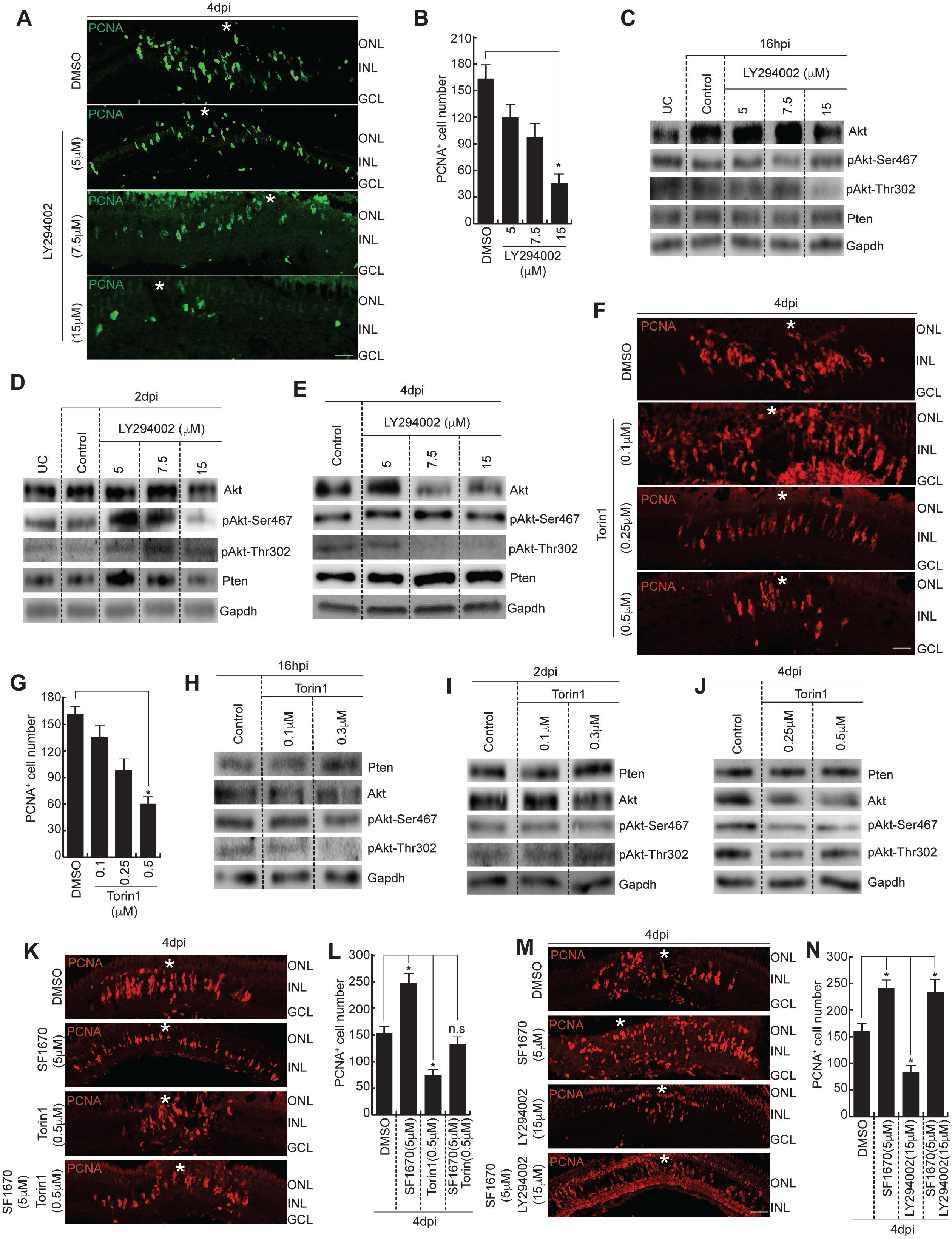
PI3K and mTORC2 are involved in the proliferation of MGPCs by phosphorylating Akt. (**A** and **B**) IF microscopy images of retinal cross-sections show a decline in the number of PCNA^+^ MGPCs with the increasing concentrations of LY294002 at 4dpi (**A**), which is quantified (**B**); *p < 0.02, n=6 biological replicates. (**C-E**) Western Blot analyses of Akt, pAkt-Ser467, pAkt-Thr302, Pten from retinal extracts collected from retinae treated with different concentrations of LY294002 at 16hpi (**C**), 2dpi (**D**) and 4dpi (**E**). (**F** and **G**) IF microscopy images of retinal cross-sections show a decline in the number of PCNA^+^ MGPCs with the increasing concentrations of Torin1 at 4dpi (**F**), which is quantified (**G**); *p < 0.004, n=6 biological replicates. (**H-J**) Western Blot analyses of Pten, Akt, pAkt-Ser467, pAkt-Thr302 from retinal extracts collected from retinae treated with different concentrations of Torin1 at 16hpi (**H**), 2dpi (**I**) and 4dpi (**J**). (**K** and **L**) IF microscopy images of retinal cross-sections show increased PCNA^+^ MGPCs with SF1670 treatment, which reduce drastically with the treatment of Torin1, while changing insignificantly in the combination of SF1670 and Torin1 as compared to the DMSO control at 4dpi (**K**), which is quantified (**L**); *p < 0.03, n=6 biological replicates. (**M** and **N**) IF microscopy images of retinal cross-sections show the reduction in the number of PCNA^+^ MGPCs with the treatment of LY294002, while an increase with SF1670 treatment, which is maintained upon the treatment with the combination of SF1670 and LY294002, as compared to the DMSO control at 4dpi (**M**), which is quantified (**N**); *p < 0.04, n=6 biological replicates. Scale bars represent 10μm in (**A, F, K, M**); the asterisk marks the injury site and GCL, ganglion cell layer; INL, inner nuclear layer; ONL, outer nuclear layer in (**A, F, K, M**); hpi, hours post injury; dpi, days post injury. Error bars represent SD; Gapdh is the loading control (**C, D, E, H, I, J**); UC-Uninjured control (**C,D**); n.s., not significant.

### Pten influences retina regeneration through Notch signaling

Notch signaling has been studied extensively in various developmental systems and is important for cell cycle progression. However, in zebrafish retina, the active notch signaling restricts the extent of proliferation of MGPCs (Conner et al., 2014; Elsaeidi et al., 2018; Mills and Goldman, 2017; Wan et al., 2012). Since there was an increase in proliferation of MGPCs in the Pten inhibited retina, we explored to find whether the Notch signaling is perturbed to facilitate this cell proliferation. We analyzed the expression levels of *her4.1* and one of its direct targets *mmp9* (Kaur et al., 2018) in the SF1670-treated retina at 4dpi (Figure 5A). We saw a drastic decline in the *her4.1*, a notch signaling effector gene, and an anticipated increase in the *mmp9* levels (Figures 5B and 5C). Furthermore, we assayed the levels of *her4.1* and *mmp9* in the double blocker experiments where mTORC2 and PI3K were blocked using Torin1 and LY294002, respectively, in combination with Pten blocker SF1670 in separate experiments.

**Figure 5:**
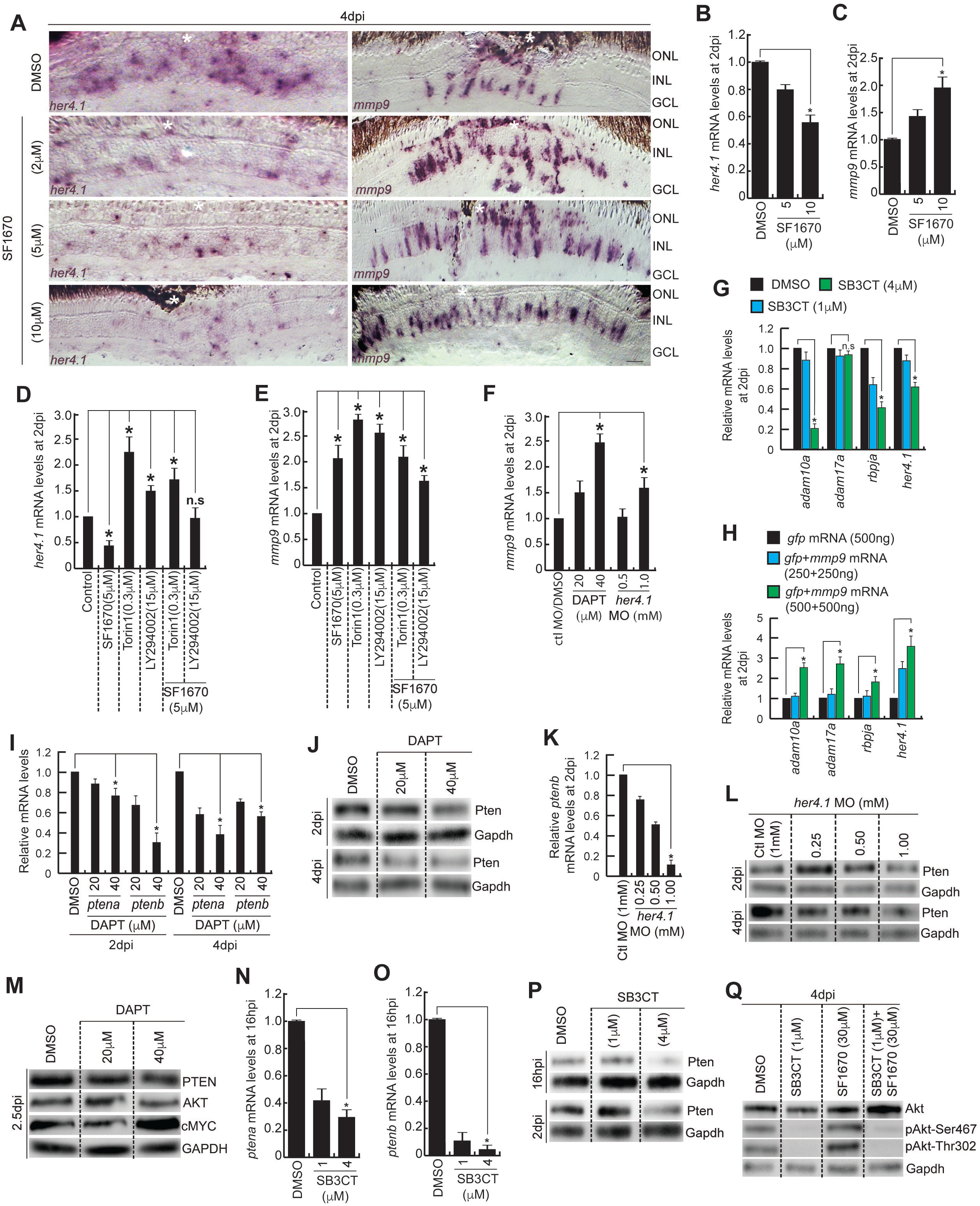
Mmp9/Notch signaling regulates Pten during retina regeneration. (**A**) Brightfield (BF) microscopy images of retinal cross-sections show the mRNA ISH of the *her4.1* and *mmp9* mRNAs in the retina treated with SF1670 at 4dpi. (**B** and **C**) The qPCR analyses of *her4.1* (**B**) and *mmp9* (**C**) mRNA levels in SF1670-treated retina at 2dpi; *p < 0.04, n=6 biological replicates. (**D** and **E**) The qPCR analyses of *her4.1* (**D**) and *mmp9* (**E**) mRNA levels in retina treated with Torin1, LY294002 alone and in combination with SF1670 at 2dpi; *p < 0.03, n=6 biological replicates. (**F**) The qPCR analyses of *mmp9* mRNA levels in DAPT-treated and *her4.1* knockdown retina at 2dpi; *p < 0.04, n=6 biological replicates. (**G**) The qPCR analyses of *adam10a*, *adam17a*, *rbpja*, *her4.1* in SB3CT-treated retina at 2dpi; *p < 0.03, n=6 biological replicates. (**H**) The qPCR analyses of *adam10a*, *adam17a*, *rbpja* and *her4.1* in *mmp9*-overexpressed retina at 2dpi; *p < 0.02, n=6 biological replicates. (**I** and **J**) The qPCR analyses of *ptena* and *ptenb* mRNA levels (**I**) and western blot analyses of Pten protein (**J**) in DAPT-treated retina at 2dpi and 4dpi; *p < 0.04, n=6 biological replicates. (**K** and **L**) The qPCR analysis of *ptenb* mRNA levels at 2dpi (**K**) and western blot analyses of Pten protein at 2dpi and 4dpi (**L**) in *her4.1* knockdown retina; *p < 0.03, n=6 biological replicates. (**M**) Western Blot analyses of PTEN, AKT, cMYC from the lysates collected from mice retina treated with different concentrations of DAPT at 2.5dpi. GAPDH is the loading control. (**N-P**) The qPCR analyses of *ptena* (**N**) and *ptenb* (**O**) mRNA levels at 16hpi and western blot analyses of Pten protein at 16hpi and 2dpi (**P**) in SB3CT-treated retina; *p < 0.02, n=6 biological replicates. (**Q**) Western Blot analyses of Akt, pAkt-Ser467, pAkt-Thr302 from retinal extracts collected from retinae treated with SB3CT alone and in combination with SF1670 at 4dpi. Gapdh is the loading control. Scale bars represent 10μm in (**A**); the asterisk marks the injury site and GCL, ganglion cell layer; INL, inner nuclear layer; ONL, outer nuclear layer in (**A**); hpi, hours post injury; dpi, days post injury. Error bars represent SD. n.s., not significant.

We saw a significant increase in *her4.1* levels where both SF1670 and Torin1 were used, and there was no appreciable effect when LY294002 was used in combination with SF1670 (Figure 5D). However, the *mmp9* levels were increasing in both sets of double blocker experiments (Figure 5E). Such a regulation might be existing because there is a selective shift in the interaction of PDK1 either with Akt or with PKC (Scheid et al., 2005) and subject to the Akt phosphorylation at its Ser467 by mTORC2. Upon Torin1 treatment, Akt is not phosphorylated, leading to the flux change of PDK1 from Akt to PKC, thus allowing Notch activation through Adams (Steinbuck and Winandy, 2018), leading to upregulation of *her4.1*, thereby keeping a check on the number of MGPCs as was seen upon Torin1 treatment in Figures 4F and 4G. There might be a feedback signal due to this decrease in the number of MGPCs, to elevate the levels of *mmp9*, which is a pro-proliferative factor, to maintain a balance in the rate of proliferation of MGPCs during retina regeneration. In this scenario, we observed an increase in the *mmp9* expression levels also due to Pten blockade, which further tried to elevate *her4.1* levels. In the combined blockade of Pten and PI3K, we observed a significant increase in the levels of *mmp9* but no change in *her4.1* levels, as compared to control. There should not have been any change in proliferation of MGPCs, levels of *her 4.1* and *mmp9*, if Pten/PI3K acted only through PDK1/Akt1/mTORC pathway. Therefore, there could be other pathway emanated from Pten, which impinges on inhibiting *mmp9* expression. Cumulatively, we saw an increase in *mmp9* levels and no significant change in *her4.1*levels, which was reflected in the extent of proliferation of MGPCs as well. The decline in *her4.1* either by blockade of notch signaling through DAPT administration or electroporation of *her4.1*-targeting MO caused an increase in the *mmp9* expression in injured retina (Figure 5F). These results suggest that the Pten blockade impinges on Notch signaling, which reduced *her4.1* that in turn accelerated *mmp9* expression.

We then explored if the increased *mmp9* in Pten-blocked retina affected the Notch signaling, which could negatively regulate the proliferation of MGPCs in the regenerating zebrafish retina (Conner et al., 2014; Elsaeidi et al., 2018; Mills and Goldman, 2017; Wan et al., 2012). The Mmp9 causes the release of the active TNFα (tumor necrosis factor-alpha) (Gearing et al., 1994; Gearing et al., 1995; Yabluchanskiy et al., 2013), which in turn upregulates NF-κB (nuclear factor kappa-light-chain-enhancer of activated B cells) (Cao et al., 1999; Feuillard et al., 1991; Fujisawa et al., 1996). The TNFα/NF-κB signaling, in turn, activates the transcription of *mmp9* (Bond et al., 1998; Nagase, 1997; Rhee et al., 2007; Sanceau et al., 2002), and also ADAM (a disintegrin and metalloproteinase) genes namely *adam17* (Wawro et al., 2019) and *adam10* (Zhu et al., 2014). Further, ADAM17 and ADAM10 are responsible for regulated intramembrane proteolysis of Notch receptors (Gibb et al., 2011; Murthy et al., 2012), which is a prerequisite for the action of the enzyme γ-secretase to produce notch intra cellular domain (NICD), which causes the transcription of effector genes such as *rbpj* and *her4.1*. Here, we blocked the Mmp9 activity using SB3CT, to assess the impact on *adam10a*, *adam17a*, *rbpja*, and *her4.1*. We saw a significant decline in all these four genes, probably because of a decrease in TNFα necessary for *adam10a*, *adam17a* expression that negatively affects Notch signaling, which brings down effector genes *rbpja* and *her4.1* at 16hpi (Figure 5G), and 2dpi (Figure EV6A). The opposite expression levels of these genes were seen when we overexpressed *mmp9* mRNA in injured retina at 2dpi (Figure 5H).

Further, the blockade of Notch signaling by DAPT treatment, which declines the *her4.1* levels, caused a reduction in *ptena* and *ptenb* mRNAs at 2dpi (Figure 5I), which is also reflected in their protein levels at both 2 and 4dpi (Figures 5J and EV6B). The reduction in Pten levels because of DAPT treatment also led to an increase in the Akt and its phosphorylated forms in the retina at 4dpi (Figures EV6C and EV6D). Similarly, the targeted knockdown of *her4.1* caused a decline in *ptenb* at 2dpi (Figure 5K), and its protein levels at both 2 and 4dpi (Figures 5L and EV6E). The *her4.1* knockdown also caused an increase in proliferation of MGPCs in the injured retina at 4dpi (Figure EV6F and EV6G). Of note, the DAPT treatment in injured mice retina did not affect Pten or Akt levels appreciably, and despite an increase in cMyc levels, only a moderate increase in EdU^+^ cells was seen in the injured mice retina at 2.5dpi (Figures 5M, EV6H, EV6I, and EV6J).

We also explored the effect of the Mmp9 blockade on Pten expression levels. We saw a dose-dependent decline in both *ptena* and *ptenb* mRNA levels in SB3CT-treated retina at 16hpi (Figures 5N and 5O), a time when *mmp9* expression and MG reprogramming was at its peak (Kaur et al., 2018; Sharma et al., 2019; Sharma et al., 2020). The SB3CT treatment caused a substantial decline in the Pten protein levels both at 16hpi and 2dpi retina (Figures 5P and EV6K). In agreement with SB3CT treatment results, the overexpression of *mmp9* mRNA caused an upregulation of *ptena* and *ptenb* mRNAs in 2dpi retina (Figure EV6L). It is important to note that the overexpression of Mmp9 could upregulate NF-κB, which is a positive regulator of PTEN expression in leukemic cells (Lee et al., 2007). This could be the reason for reduced Pten levels in the SB3CT-treated retina. We then explored the levels of Akt and its phosphorylated forms in SF1670-treated retina in the absence of Mmp9 activity using SB3CT. Interestingly, we saw a dramatic decline in the phosphorylated forms of Akt despite an increase in Akt levels (Figures 5Q and EV6M). These results support the view that Mmp9 activity is indispensable for the phosphorylation of Akt and thus proliferative response observed with Pten blockade. Taken together, these observations suggest that Pten affects proliferation of MGPCs through Notch signaling, which in turn is regulated by Mmp9.

### Regulation of Pten by pathways other than Mmp9/Notch signaling influences the proliferation of MGPCs

We explored whether the increase in the number of MGPCs seen in Pten blocked condition is mediated through Mmp9. To investigate this, we did a double blocker experiment in which injured retina was treated with both SF1670 and SB3CT drugs. We observed that separate treatments of SF1670 and SB3CT caused an anticipated increase and decrease in the number of MGPCs, respectively, in the retina at 4dpi (Figures 6A and 6B). However, when both SF1670 and SB3CT were administered, the increase in MGPCs because of SF1670 was nullified, and on the contrary, the observed decrease in cell proliferation in SB3CT-treated retina got increased to a level close to control injured retina at 4dpi (Figures 6A and 6B). Even the selective blockade of *mmp9* using MO also had a similar effect on the proliferation of MGPCs (Figures 6C and 6D). These results confirm that the observed increase in MGPCs number in Pten-blocked retina requires Mmp9 activity, and conversely, the block in cell proliferation because of inactive Mmp9 could be alleviated by Pten inhibition, suggesting the existence of different cellular pathways in the absence of Pten to regulate the proliferation of MGPCs.

**Figure 6:**
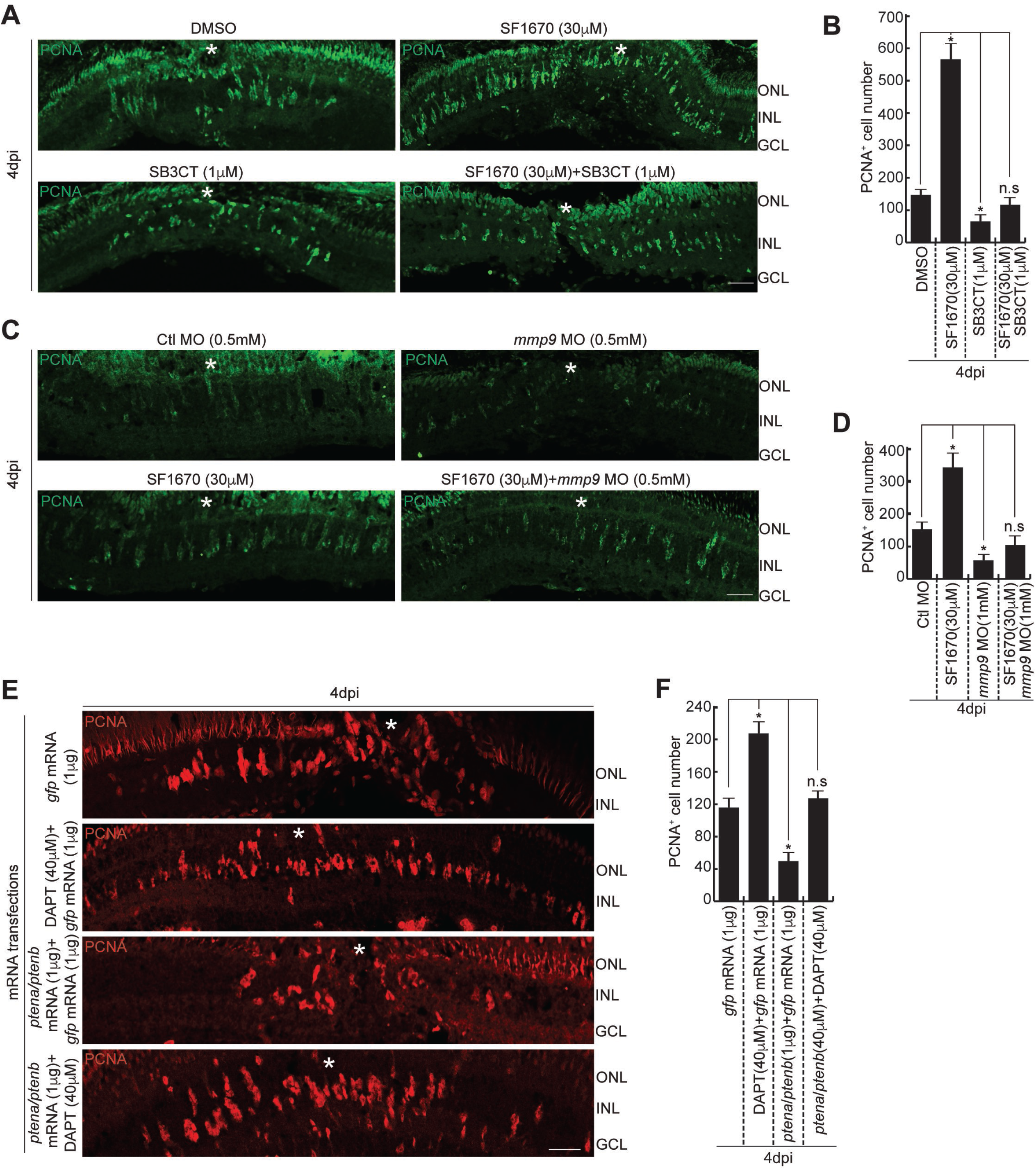
Involvement of different pathways to influence Pten function during retina regeneration. (**A** and **B**) IF microscopy images of retinal cross-sections show an increase in the number of PCNA^+^ MGPCs with SF1670 treatment, which reduces drastically with the treatment of SB3CT, while the number is elevated close to the DMSO control in the combination of SF1670 and SB3CT at 4dpi (**A**), which is quantified (**B**); *p < 0.04, n=6 biological replicates. (**C** and **D**) IF microscopy images of retinal cross-sections show an increase in the PCNA^+^ MGPCs with SF1670 treatment, which decreases with the *mmp9* knockdown, while this number is elevated close to the DMSO control in the combination of SF1670 and *mmp9* MO in 4dpi retina (**C**), which is quantified (**D**); *p < 0.04, n=6 biological replicates. (**E** and **F**) IF microscopy images of retinal cross-sections show a significant increase in the number of PCNA^+^ MGPCs with DAPT treatment, which decreases with the *ptena/ptenb* overexpression and gets close to the DMSO control in the combination of DAPT and *ptena*+*ptenb* mRNAs-treated retina at 4dpi (**E**), which is quantified (**F**); *p < 0.03, n=6 biological replicates. Scale bars represent 10μm in (**A, C, E**); the asterisk marks the injury site and GCL, ganglion cell layer; INL, inner nuclear layer; ONL, outer nuclear layer in (**A, C, E**); dpi, days post injury. Error bars represent SD. n.s., not significant.

In a similar scenario, the DAPT-mediated block of Notch signaling causing an enhanced proliferation of MGPCs was analyzed in Pten overexpressed background. The purpose of this experiment was to assess if the role of Pten during retina regeneration was influenced by Notch signaling. Overexpression of Pten significantly reduced the number of MGPCs in the injured retina, which could also nullify the effect of DAPT-treatment (Figures 6E and 6F). These observations suggest that reduced Pten levels are a necessity for the induction of MGPCs in both normal and DAPT-treated retina. The blockade of Notch signaling, causing an increase in the proliferation of MGPCs, should be imperatively linked to Pten reduction as shown (Figures 5I, 5J and EV6B). These findings also suggest that Pten could influence the proliferation of MGPCs through means other than the Notch signaling.

### Regulation of *pten* expression during retina regeneration

We explored how the *pten* expression was regulated during retina regeneration. The *pten* gene is known to be regulated by a plethora of regulatory molecules, and hence finding a plausible regulatory mechanism of *pten* was rather challenging. It is to be noted that the Pten levels should decline after an injury to facilitate the proliferation of MGPCs, which should then be restored back to prevent unwanted, persistent proliferation. To analyze this, we explored the promoter elements of *ptena* and *ptenb* genes. We found a number of Mycb-binding sites on the promoter sequence of both *ptena* and *ptenb* genes. Mycb is reported to act as a transcriptional activator or repressor depending upon its collaborating partner (Herkert and Eilers, 2010). During zebrafish retina regeneration, Mycb is seen to collaborate with Hdac1 to regulate *lin28a* expression (Mitra et al., 2019). Both Mycb (Mitra et al., 2019; Ramachandran et al., 2010) and Hdac1 (Mitra et al., 2018) are induced soon after injury in zebrafish retina which physically collaborates to occupy *lin28a* promoter to cause its downregulation (Mitra et al., 2019). In these lines of thought, we explored if the *ptena* and *ptenb* levels are affected when Mycb or Hdac1 are inhibited. We used 10058-F4 (Huang et al., 2006; Lin et al., 2007; Wang et al., 2007; Yin et al., 2003), and Trichostatin A (TSA) (Bolden et al., 2006; Xu et al., 2007) which block Myc-Max interaction and Hdac1, respectively. We saw a significant increase in the *ptena* and *ptenb* mRNAs because of the 10058-F4 drug treatment in the retina at 2dpi (Figures 7A and 7B), which was also reflected in the protein levels at 4dpi (Figure 7C). Similar results were also seen with TSA treatment (Figures 7D-7F), which suggests the possibility of repressive gene regulation of *ptena* and *ptenb* mediated by Mycb-Hdac1 complex. We then performed chromatin immunoprecipitation (ChIP) using Mycb and Hdac1-specific antibodies separately to see whether the Mycb binding sites of *ptena* and *ptenb* gene promoters were occupied by Mycb-Hdac1 complex. Interestingly, we saw the co-occupancy of Mycb and Hdac1 on all 3 sites present on the *ptena* and *ptenb* promoters, which confirmed the repression of *pten* genes by Mycb-Hdac1 complex (Figure 7G). We further validated the ChIP data by adopting *1016tuba1a*:GFP transgenic zebrafish. We sorted the GFP^+^ and GFP^-^cell population at 4dpi before performing the ChIP using anti-Mycb and anti-Hdac1 antibodies, separately. The ChIP qPCR results also showed that Mycb and Hdac1 both bind on the Mycb-binding sites of *ptena*/*ptenb* promoters majorly in GFP^+^ cells than in the GFP^−^ ones (Figure 7H and 7I). These findings support the view that Mycb-Hdac1 complex negatively regulates Pten expression during retina regeneration.

**Figure 7:**
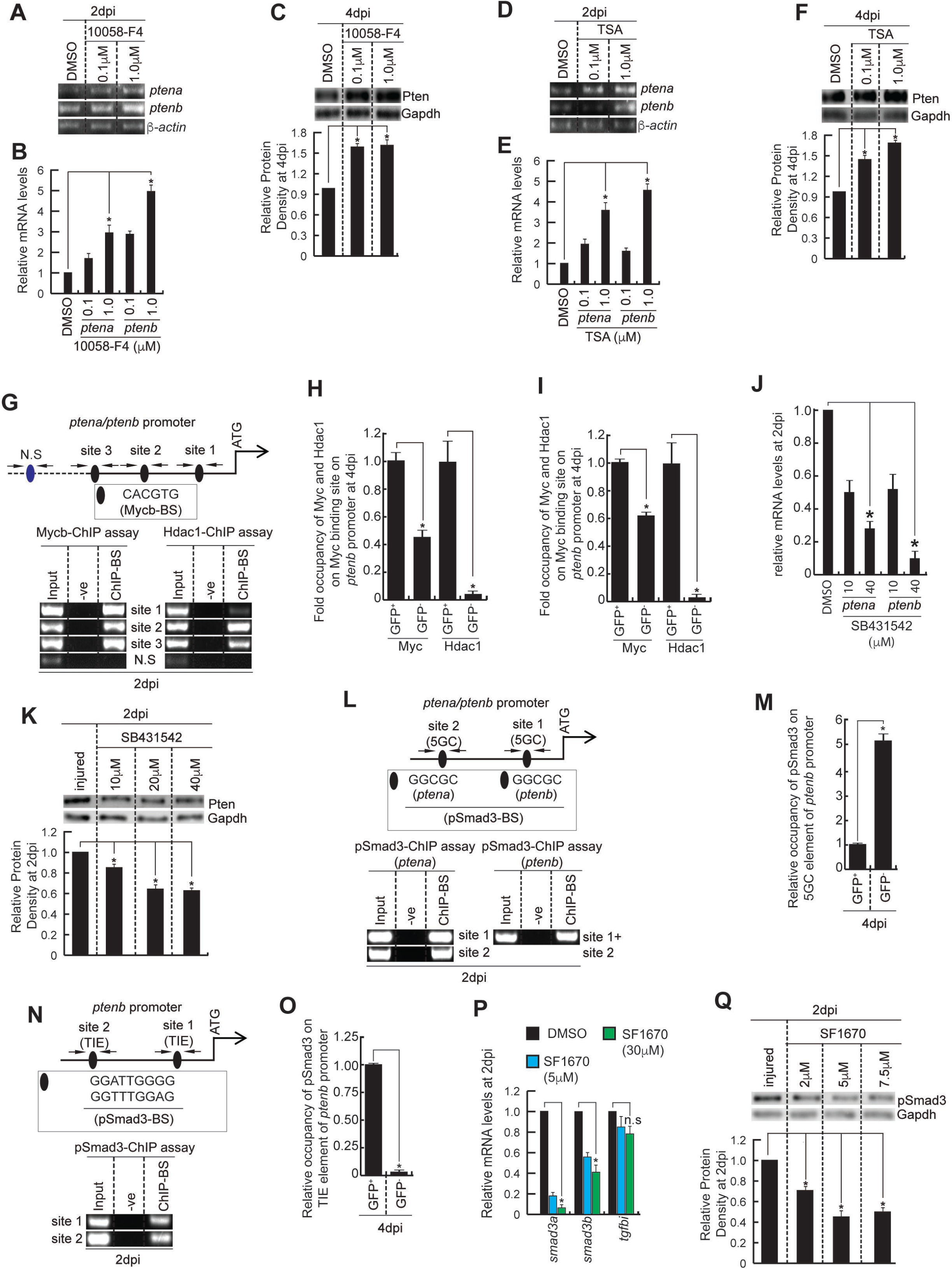
Fine-tuned regulation of Pten during retina regeneration. (**A** and **B**) The RT-PCR (**A**) and qPCR (**B**) analyses of *ptena* and *ptenb* mRNA levels in 10058-F4-treated retina at 2dpi; *p < 0.025, n=6 biological replicates. (**C**) Western blot analysis (above) of Pten protein in 10058-F4-treated retina at 4dpi, quantified by densitometry (below); *p < 0.001; n=6 biological replicates. (**D** and **E**) The RT-PCR (**D**) and qPCR (**E**) analyses of *ptena* and *ptenb* mRNA levels in TSA-treated retina at 2dpi; *p < 0.005, n=6 biological replicates. (**F**) Western blot analysis (above) of Pten protein in TSA-treated retina at 4dpi, quantified by densitometry (below); *p < 0.001; n=6 biological replicates. (**G**) The *ptena/ptenb* promoter schematic reveals the Mycb-binding sites (BS) (upper), and the retinal ChIP assays confirm the physical binding of Mycb along with Hdac1 to those sites (lower), in 2dpi retina. N.S marks the negative control, and capital letters mark putative Mycb-BS. (**H**) The ChIP-qPCR analysis shows the relative occupancy of Mycb and Hdac1 on putative Mycb-BS on *ptenb* promoter in GFP^+^ and GFP^−^ cells from *1016tuba1a*:GFP transgenic retina at 4dpi; *p < 0.001, pooled sample from 20 retina at 4dpi. **I**) The ChIP-qPCR analysis shows the relative occupancy of Mycb and Hdac1 on putative Mycb-BS on *ptena* promoter in GFP^+^ and GFP^−^ cells from *1016tuba1a*:GFP transgenic retina at 4dpi; *p < 0.001, pooled sample from 20 retina at 4dpi. (**J)** The qPCR analyses of *ptena* and *ptenb* levels in SB431542-treated retina at 2dpi; *p < 0.03, n=6 biological replicates. (**K**) Western blot analysis of Pten in SB431542-treated retina at 2dpi (above), quantified by densitometry (below); *p < 0.001; n=6 biological replicates. (**L**) The *ptena/ptenb* promoter schematic reveals the typical 5GC sites (upper) and the retinal ChIP assays confirm the physical binding of pSmad3 at the 5GC sites (lower) in 2dpi retina. Capital letters mark 5GC sequence. (**M**) The ChIP-qPCR analysis shows the relative occupancy of pSmad3 on 5GC element of *ptenb* promoter in GFP^+^ and GFP^−^ cells from *1016tuba1a*:GFP transgenic retina at 4dpi; *p < 0.001, pooled sample from 20 retina at 4dpi. (**N**) The *ptenb* promoter schematic reveals the typical TIE sequence (upper) and the retinal ChIP assay confirms the physical binding of pSmad3 at the TIE sites (lower) in 2dpi retina. Capital letters mark the TIE sequences. (**O**) The ChIP-qPCR analysis shows the relative occupancy of pSmad3 on TIE sequence of *ptenb* promoter in GFP^+^ and GFP^−^ cells from *1016tuba1a*:GFP transgenic retina at 4dpi; *p < 0.001, pooled sample from 20 retina at 4dpi. (**P**) The qPCR analyses of Tgf-β signaling reporter genes *smad3a*, *smad3b* and *tgfbi* mRNA levels in SF1670-treated retina at 2dpi; *p < 0.03, n=4 biological replicates. (**Q**) Western blot analysis of pSmad3 in SF1670-treated retina at 2dpi (above), quantified by densitometry (below); *p < 0.001; n=6 biological replicates. dpi, days post injury; Arrows mark ChIP primers in (**G**), (**L**) and (**N**); Error bars represent SD; Gapdh is the loading control (**C, F, K, Q**); Ctl MO-Control morpholino (**C**); n.s., not significant.

We further explored if there could exist a fine-tuned mechanism of *pten* regulation. At first, we explored the effect of Tgf-β signaling, which has a pro-proliferative role during zebrafish retina regeneration (Sharma et al., 2020), on Pten. For this, we treated the retina with the drug SB431542, which blocks Tgf-β signaling (Halder et al., 2005; Sharma et al., 2020), and checked its effect on *pten* gene expression. We saw a dose-dependent reduction in both *ptena* and *ptenb* genes (Figure 7J), as well as protein expression (Figure 7K) in SB431542-treated retina at 2dpi. Promoter analysis of *ptena* and *ptenb* genes revealed the presence of typical 5GC sites, GGC(GC)/CG, on which pSmad3 binds to activate gene expression (Martin-Malpartida et al., 2017). We further confirmed the occupancy of these 5GC elements by pSmad3 by ChIP assay using pSmad3-specific antibodies at 2dpi (Figure 7L). We further confirmed the ChIP data by making use of *1016tuba1a*:GFP transgenic zebrafish. We sorted the GFP^+^ and GFP^−^ cell population at 4dpi before performing the ChIP using anti-pSmad3 antibody. The ChIP PCR is quantified by qPCR (Figure 7M). The results support the previous observation that pSmad3 binds effectively on the 5GC elements of *ptenb* promoter in GFP^−^ cells where the Pten expresses abundantly. These findings support the view that Tgf-β signaling positively regulates Pten expression during retina regeneration. However, activation of Tgf-β signaling is always shown to facilitate the proliferation of MGPCs, which seemed like a conundrum. For this, we closely looked into the promoters of *ptena* and *ptenb* and found that there is a *bona fide* Tgf-β inhibitory element (TIE), a unique DNA sequence essential for the transcriptional repression events mediated by TGF-β1 (Kerr et al., 1990). We saw a TIE sequence on the *ptenb* gene promoter, which is found to be occupied by pSmad3, as revealed in a ChIP assay using a pSmad3-specific antibody (Figure 7N). Again, we made use of the *1016tuba1a*:GFP transgenic zebrafish to validate the differential occupancy of pSmad3 on TIE sequence of *ptenb* promoter in GFP^+^ and GFP^−^ cell population. The ChIP assay followed by qPCR revealed that the pSmad3 occupied TIE sequences more efficiently in the GFP^+^ cells (Figure 7O). This observation explains the reduced *pten* expression in GFP^+^ MGPCs. However, the pSmad3 could occupy the TIE only when Fos, a product of *the cfos* gene, was present as part of the binding complex (Kerr et al., 1990). The *cfos* gene gets rapidly induced in zebrafish retina (Figure EV6N and EV6O), and is associated with the proliferating MGPCs (Figure EV6P) where the *pten* expression stays downregulated. These observations suggest that Tgf-β signaling could positively regulate the Pten expression through 5GC, and negatively in the presence of Fos through TIE.

Further, we checked if Pten has any regulatory role on Tgf-β signaling. Upon Pten-blockade by SF1670, we saw a downregulation of components of Tgf-β signaling, namely *smad3a*, *smad3b*, and an insignificant change in the levels of effector gene *tgfbi* in the retina at 2dpi (Figure 7P). The pSmad3 protein levels were also seen to decrease with SF1670 treatment in 2dpi retina (Figure 7Q). The SF1670 treatment caused downregulation of *her4.1*, which is a reporter of active Notch signaling mediated by NICD. Overexpression of NICD is known to enhance Tgf-β signaling through pSmad (Misra et al., 2014). This study could explain the decreased *smad3a*, *smad3b*, and *tgfbi* in SF1670-treated retina. These findings suggest that Pten positively regulates its own expression through Tgf-β signaling during retina regeneration. This type of dual *pten* regulation and Pten’s influence on Tgf-β signaling ensures the rapid return of Pten to avoid the undesirable hyper-proliferation of MGPCs during zebrafish retina regeneration.

## Discussion

In this study, we explored the significance of the tumor suppressor Pten and the dynamics of its regulation during zebrafish retina regeneration. Pten is the second most mutated tumor suppressor after p53 to cause cancer, which suggests the potential of a Pten-mediated pathway in retaining cellular homeostasis. Despite the knowledge about the Pten-PI3K-Akt-mTOR pathway in many cell proliferative phases such as development and cancer, little is known about its significance during zebrafish retina regeneration. In zebrafish, the successful retina regeneration is the result of MG reprogramming that pave the way to the formation of a proliferating population of MGPCs, which give rise to almost all retinal cell-types. These MGPCs express several regeneration-specific genes and several pluripotency inducing factors, yet do not lead to undesirable proliferation which could potentially cause cancer. In zebrafish, however, the *ptenb* loss of function mutation has been reported to cause spontaneous retinal tumors, which suggested the potential of retinal cells to divert into cancerous conditions (Faucherre et al., 2008). We decided to explore the importance of *pten* gene regulation and the biological functions of Pten during zebrafish retina regeneration. The findings from this study are summarized and organized in a model as detailed (Figure EV7).

The expression pattern of both *pten* genes in the injured retina agrees with the reported role of Pten as a tumor suppressor (Dahia, 2000; Simpson and Parsons, 2001). The selective downregulation of Pten in MGPCs seemed a remarkable strategy of the cells to allow the pro-proliferative PI3K/Akt/mTOR pathway to ensue. The blockade of Pten function either by gene downregulation or pharmacological inhibition showed an accelerated proliferative response of MGPCs while retaining their ability to differentiate into various retinal cell types. Absence of Pten function is one of the major causes of uncontrolled cell proliferation in cancer, which makes it one of the most important tumor suppressors similar to p53 in maintaining cellular homeostasis (Ming and He, 2012; Parsons and Simpson, 2003; Simpson and Parsons, 2001; Sulis and Parsons, 2003). PTEN is also involved in a co-operative interrelationship with p53 as part of its functions as a tumor suppressor (Baron et al., 2006; Mayo and Donner, 2002; Nakanishi et al., 2014; Okumura et al., 2011). In zebrafish, the downregulation of Pten is observed in MGPCs, and its downregulation promotes cell proliferation, while its overexpression reduced proliferating cell populations. These attributes strongly support the involvement of Pten in retinal homeostasis as a suppressor of cell proliferation. Further, the modulation of Akt in its wild-type form or a phosphomimetic one in the zebrafish retina also supported its role to cause accelerated cell proliferation. The activation of Akt by its phosphorylation at T-302 and S-467 by PI3K and mTORC2 respectively enables its cell proliferative function that is reversed upon blockade of PI3K and mTORC2. A recent study also demonstrated that the post-injury retinal inflammation could accelerate mTOR, which is necessary prior to regenerative response (Zhang et al., 2020).

Interestingly, our double blocker experiments to block mTORC2 along with Pten inhibition demonstrated that the enhanced number of MGPCs seen with SF1670 indeed is associated with the phosphorylated form of Akt at S-467. In the absence of phosphorylation at S-467 of Akt, the Pten blockade was inadequate to cause an enhancement of the number of MGPCs. On the other hand, in the double blocker experiments with PI3K and Pten inhibition, there existed a similar number of MGPCs as found in Pten inhibition alone, suggesting that the effect by the blockade of phosphorylation at T-302 of Akt gets nullified when Pten inhibition is accompanied with it. Earlier studies have reported that the phosphorylation of T-308 by phosphoinositide-dependent kinase 1 (PDK1), an enzyme activated by PI3K, is facilitated if the substrate is S-473 phosphorylated Akt1 than its wild-type form (Huang et al., 2008; Sarbassov et al., 2005). Such a scenario could exist during retina regeneration, which could explain these results. Nonetheless, the absence of a functional phosphorylated form of Akt was not enough to block the induction of MGPCs, which suggested the possible existence of other pro-proliferative pathways during retina regeneration. Based on this assumption, we explored if Pten blockade was involved in the gene expressions other than those mediated through Akt pathway. We saw a decline in the expression of *her4.1* in Pten blocked retina and an accelerated *mmp9* expression, that pave the way for upregulation of *adam10a* and *adam17a* whose activity has implications on a plethora of biological functions including cell proliferation (Huovila et al., 2005; Rocks et al., 2008). We also showed that the blockade of Notch signaling or *her4.1* knockdown caused a decline in Pten levels, which also could explain the accelerated proliferation of MGPCs in regenerating retina, which is also seen in mice retina. Also, the blockade of Mmp9 could repress Pten levels, but could not cause an increase in MGPCs as the Mmp9 itself is an indispensable molecule in retina regeneration as reported (Kaur et al., 2018; Sharma et al., 2019; Sharma et al., 2020). However, a recent study of retina regeneration after photobleaching (Silva et al., 2020) showed anti-proliferative role of Mmp9 during the photoreceptor regeneration. The discrepancy in the role of Mmp9 could be because of the mode of injury. The mechanical injury is focal and uniform across the retinal layers, whereas the photobleaching damages preferentially the photoreceptors.

Our study, while showing the importance of Pten downregulation in MGPCs as a necessary requirement, also showed the importance of Mmp9 and Notch signaling, which work in concert with Pten/PI3K/Akt axis. We performed a few double blocker experiments in which Pten was blocked along with Mmp9 inhibition to assess the existence of any hierarchical or parallel pathways that influence the number of MGPCs in the regenerating retina. Mmp9 blockade is known to adversely affect the formation of MGPCs in zebrafish retina (Kaur et al., 2018; Sharma et al., 2019; Sharma et al., 2020). However, along with Pten blockade, the number of MGPCs that are formed is arguably equal to that in injured control retina, where the variation in number was statistically insignificant. Similarly, the accelerated MGPCs seen in Notch signaling inhibited retina of zebrafish (Conner et al., 2014; Elsaeidi et al., 2018; Kaur et al., 2018; Mills and Goldman, 2017; Mitra et al., 2018; Mitra et al., 2019; Sharma et al., 2020; Wan et al., 2012) could be nullified using Pten overexpression. Conversely, it is also interesting to note that the anti-proliferative effect of the Pten overexpression was alleviated in combination with DAPT treatment. These observations suggest the possible existence of different pathways to Pten/Akt that govern the proliferation of MGPCs, which could impinge on each other to ensure the adequate number of MGPCs in the regenerating retina.

Further, we were intrigued to find the fine balancing of the Pten which seemed essential for the optimal regenerative response. In this study, we showed that over or under activation of Pten had a dose-dependent link to the proliferation of MGPCs. This observation made us to explore the mechanisms through which the Pten could be regulated. There are several studies which discuss the fine regulation of Pten expression in various biological systems (Bermudez Brito et al., 2015). However, here we explored the *pten* regulations through already reported essential pathways for normal regeneration. One of such mechanisms was through the Mycb-Hdac1 collaboration to repress the *pten* gene. We showed that the Hdac1 in collaboration with Mycb occupied the E-boxes present on *pten* promoter for a repressive cause, and the inhibition of either Mycb or Hdac1 accelerated the *pten* expression, with a negative effect on the number of MGPCs as reported (Mitra et al., 2018; Mitra et al., 2019). Similarly, another very important signaling pathway, the Tgf-β signaling, which is known to be essential for normal regenerative response in zebrafish retina (Sharma et al., 2020) regulated the *pten* in two contrasting ways. At first, we inhibited the Tgf-β signaling to assess its effect on *pten* expression and found that Tgf-β signaling and *pten* expression are positively correlated. We could confirm this through ChIP assay of pSmad3, the effector of Tgf-β signaling, on *pten* promoter at 5GC sites (Martin-Malpartida et al., 2017). This observation apparently seemed a conundrum as we knew that Tgf-β signaling played a pro-proliferative role, and Pten does an anti-proliferative role. Interestingly, we could also find unique TIE sequences which could be a means of gene repression through binding of pSmad3 along with other immediate early induced factors such as Fos to inhibit Pten expression. In other words, although Tgf-β signaling could upregulate Pten expression through 5GC elements, the TIE sequences could enable the downregulation in selected cells such as MGPCs. The absence of Fos at later stages of retina regeneration, could bring up the Pten levels as an essential step to exit cell cycle. Pten inhibition caused a downregulation of Tgf-β signaling components, which in turn lowers *pten* levels. In other words, Pten could positively regulate its own expression through Tgf-β signaling. Thus, the robust regulation of Pten through various transcription factors and cellular signaling events ensures induction of adequate MGPCs during zebrafish retina regeneration.

## Materials and Methods

Experiments are performed as described previously (Fausett and Goldman, 2006; Sharma and Ramachandran, 2019). Further details and an outline of resources used in this work can be found in Supplemental Experimental Procedures.

## Acknowledgments

S.G acknowledges her support from the ICMR for Senior Research Fellowship. P.S acknowledges postdoctoral fellowship support from Wellcome Trust/DBT India Alliance, IISER Mohali and DST India. M.C acknowledges her financial support from the IISER Mohali. S.K acknowledge DBT for Senior Research Fellowship. V.K.V acknowledges support from IISER Mohali. This work was supported by the Wellcome Trust/DBT India Alliance Intermediate Fellowship awarded to Rajesh Ramachandran (IA/I/12/2/500630). R.R also acknowledges research funding from Science Education and Research Board SERB, DST, India (EMR/2017/001816), DBT India (BT/PR17912/ MED/31/336/2016) and support from IISER Mohali.

## Author Contributions

R.R. conceived the study and designed experiments. S.G. performed the majority of experiments. M.C. contributed to few of the western blotting assays, immunostainings and mRNA *in-situ* hybridisation experiments and helped during FACS experiment. P.S. performed the mice experiments and Fluorescence *in situ* hybridisation experiments. V.K.V contributed to *cfos* gene time-course data. S.K. provided the TOPO-*ptenb* CDS clone. R.R. wrote the manuscript with critical inputs from S.G., P.S.,M.C and S.P.

## Expanded View Figure Legends

**Figure EV1: Spatial expression pattern of *ptena* and *ptenb* genes in the retina.**

(**A**) CLUSTALW analysis shows the amino acid sequence alignment between zebrafish Ptena and Ptenb proteins. (**B**) Fluorescence *in-situ* hybridisation (FISH) and Immunofluorescence (IF) microscopy images of retinal cross-sections reveal the expression of *ptena* and *ptenb* mRNA majorly in the neighbouring cells of BrdU^+^ MGPCs at 2dpi, 4dpi and 6dpi. (**C**) Brightfield (BF) and IF microscopy images of broader span of retinal cross-sections show the mRNA *in-situ* hybridisation which reveals the expression of *ptena* and *ptenb* mRNA majorly in the neighbouring cells of BrdU^+^ MGPCs at 2dpi, 4dpi and 6dpi. (**D**) IF microscopy images of a retinal cross-section show the cytoplasmic and nuclear localised Pten in the neighbouring cells of MGPCs while being almost absent from the BrdU^+^ MGPCs at 4dpi; DAPI is the nuclear counterstain. (**E**) IF microscopy images of a retinal cross-section show the expression of Pten in the Müller Glia cells of an uninjured retina of *gfap*:GFP transgenic fish. (**F** and **G**) The qPCR analyses of *ptena* (**F**) and *ptenb* (**G**) mRNA from GFP^−^ cells present in rest of the retina compared to the GFP^+^ MGPCs in the retina of *1016tuba1a*:GFP transgenic fish at 4dpi; *p < 0.04, n=12 biological replicates. Scale bars represent 10μm in (**B, C, D, E**); the asterisk marks the injury site and GCL, ganglion cell layer; INL, inner nuclear layer; ONL, outer nuclear layer in (**B, C, D, E**); dpi, days post injury; white arrowheads mark BrdU^+^/*ptena*^−^ (**B** and **C**), BrdU^+^/*ptenb*^−^ (**B** and **C**), BrdU^+^/Pten^−^ (**D**) cells; white arrows mark *ptena*^+^/BrdU^−^ (**B** and **C**), *ptenb*^+^/BrdU^−^ (**B** and **C**), Pten^+^/BrdU^−^ (**D**) cells; Error bars represent SD.

**Figure EV2: Rescue of the effect of *pten* knockdown by *pten* mRNA overexpression.**

(**A** and **B**) IF microscopy images of retinal cross-sections show an increase in the number of PCNA^+^ MGPCs in the combined knockdown of *ptena* and *ptenb* in retina at 4dpi (**A**), which is quantified (**B**); *p < 0.01, n=6 biological replicates. (**C** and **D**) IF microscopy images of retinal cross-sections show the rescue of *ptena* MO effect by the transfection of *ptena* MO-binding site mutated mRNA in retina at 4dpi (**C**), which is quantified (**D**); *p < 0.02, n=6 biological replicates. (**E** and **F**) IF microscopy images of retinal cross-sections show the rescue of *ptenb* MO effect by the transfection of *ptenb* MO-binding site mutated mRNA in retina at 4dpi (**E**), which is quantified (**F**); *p < 0.015, n=6 biological replicates. (**G** and **H**) IF microscopy images of retinal cross-sections show a decrease in the number of BrdU^+^ MGPCs upon the combined overexpression of *ptena* and *ptenb* mRNA in retina at 4dpi (**G**), which is quantified (**H**); *p < 0.04, n=6 biological replicates. (**I**) IF microscopy images of retinal cross-sections show a concentration dependent increase in the GFP expression in *1016tuba1a*:GFP transgenic zebrafish line upon SF1670 treatment at 4dpi; n=6 biological replicates. (**J**) IF microscopy images of a retinal cross-section show that the proliferative response marked by PCNA^+^ MGPCs is absent in the uninjured retina treated with SF1670; n=6 biological replicates. Scale bars represent 10μm in (**A, C, E, G, I, J**); the asterisk marks the injury site and GCL, ganglion cell layer; INL, inner nuclear layer; ONL, outer nuclear layer in (**A, C, E, G, I, J**); dpi, days post injury; Ctl MO-Control morpholino (**A-F**), *ptena/ptenb*-MM MO is mismatch MO (**C-F**); Error bars represent SD. n.s., not significant.

**Figure EV3: Effect of Pten blockade by dipping in SF1670 on the proliferation of MGPCs, effect of *pten* downregulation and blockade on β-catenin expression and the number of mitotically active cells, TUNEL assay, Retinal cell-type staining.**

(**A** and **B**) IF microscopy images of retinal cross-sections show an increase in the number of PCNA^+^ MGPCs upon Pten blockade by dipping the fish in SF1670 (10μM) during different phases of retina regeneration (**A**), which is quantified (**B**); *p < 0.01, n=6 biological replicates. (**C**) IF microscopy images of retinal cross-sections show an insignificant number of TUNEL^+^ cells in *ptena* and *ptenb* knockdown and in SF1670 treatment as compared to the control retina at 2dpi; n=6 biological replicates. (**D** and **E**) IF microscopy images of retinal cross-sections show an increase in the number of mitotically active pH3^+^ cells upon *ptena* and *ptenb* knockdown at 4dpi (**D**), which is quantified (**E**); *p < 0.02, n=6 biological replicates. (**F** and **G**) IF microscopy images of retinal cross-sections show an increase in the expression of β-catenin in *ptenb* knockdown (**F**) and SF1670 treatment (**G**) at 4dpi; n=6 biological replicates. (**H**) An experimental timeline that describes drug delivery at the time of injury, BrdU exposure for 4hrs for 2-5dpi, followed by harvesting at 30dpi. (**I**) IF microscopy images of retinal cross-sections show the increased number of BrdU^+^ cells in SF1670-treated retina at 30dpi which make various retinal cell types, where Müller Glia are marked by Glutamine Synthetase (GS), Bipolar cells are marked by PKC-β1, Amacrine cells are marked by HuC/D. Scale bars represent 10μm in (**A, C, D, F, G, I**); the asterisk marks the injury site; GCL, ganglion cell layer; INL, inner nuclear layer; ONL, outer nuclear layer in (**A, C, D, F, G, I**); dpi, days post injury; white arrowheads mark BrdU^+^/GS^+^ (**I**), BrdU^+^/PKC-β1^+^ (**I**) and BrdU^+^/HuC/D^+^ (**I**) cells; Ctl MO-Control morpholino (**C-F**); Error bars represent SD.

**Figure EV4: Akt activation mediated by Pten blockade, Regulation of Pten and Akt by NMDA-mediated retinal injury, Regulation of proliferation of MGPCs and Akt activation by Rapamycin-mediated mTORC1 blockade.**

(**A-D**) Western Blot analyses of Pten, Akt, pAkt-Ser467, pAkt-Thr302 from retinal extracts prepared from retinae injected with different concentrations of SF1670 at 16hpi (**A**) and 2dpi (**C**), quantified by densitometry (**B** and **D**); *p < 0.04; n=6 biological replicates. (**E**) Whole retina IF microscopy image shows the proliferative response in the entire retina, as marked by BrdU^+^ MGPCs, upon panretinal NMDA-mediated injury at 4dpi. (**F**) IF microscopy images of a retinal cross-section show the Pten expression in the neighbouring cells of MGPCs while being almost absent from the BrdU^+^ MGPCs in NMDA-mediated injured retina at 4dpi; DAPI is the nuclear counterstain. (**G** and **H**) Western Blot analyses of Pten, Akt, pAkt-Ser467, pAkt-Thr302 from retinal lysates prepared from needle poke-injured and NMDA-mediated-injured retinae at 4dpi, compared with the uninjured retina (**G**), quantified by densitometry (**H**); *p < 0.001; n=6 biological replicates. (**I** and **J**) IF microscopy images of retinal cross-sections show a decline in the number of PCNA^+^ MGPCs with the increasing concentrations of Rapamycin at 4dpi (**I**), which is quantified (**J**); *p < 0.035, n=6 biological replicates. (**K** and **L**) Western Blot analyses of Akt, pAkt-Ser467, pAkt-Thr302 from retinal extracts prepared from retinae treated with different concentrations of Rapamycin at 4dpi (**K**), quantified by densitometry (**L**); *p < 0.001; n=6 biological replicates. Scale bars represent 10μm in (**E, F, I**); the asterisk marks the injury site; GCL, ganglion cell layer; INL, inner nuclear layer; ONL, outer nuclear layer in (**E, F, I**); hpi, hours post injury; dpi, days post injury. White arrowheads mark PCNA^+^/Pten^−^ (**F**); white arrows mark Pten^+^/PCNA^−^ (**F**). Error bars represent SD; Gapdh is the loading control (**A, C, G, K**); UC-Uninjured control (**A-D,G,H**); n.s., not significant.

**Figure EV5: Rescue of the effect of *akt1* knockdown. Regulation of Pten, Akt and its phosphorylated forms upon Akt activation and PI3K and mTORC2 blockade.**

(**A** and **B**) IF microscopy images of retinal cross-sections show the rescue of *akt1* MO effect by the transfection of *akt1* MO-binding site mutated mRNA in retina at 4dpi (**A**), which is quantified (**B**); *p < 0.02, n=6 biological replicates. (**C** and **D**) Western Blot analyses of Akt, pAkt-Ser467, pAkt-Thr302 from retinal lysates prepared from retinae transfected with different concentrations of *akt1* mRNA with neutral mutation and with phosphomimetic mutation, as compared to the wild-type *akt1* mRNA transfected retina at 4dpi (**C**), quantified by densitometry (**D**); *p < 0.04; n=6 biological replicates. Gapdh is the loading control. (**E-J**) Densitometry plots for Western blots (corresponding to **Figure 4C-E, H-J**) show the regulation of Pten, Akt, pAkt-Ser467, pAkt-Thr302 in LY294002-treated and Torin1-treated retinae at 16hpi (**E** and **H**), 2dpi (**F** and **I**) and 4dpi (**G** and **J**); *p < 0.01; n=6 biological replicates. (**K**) IF microscopy images of retinal cross-sections show the TUNEL^+^ cells in LY294002 and Torin1 treatment as compared to the control retina at 4dpi; n=6 biological replicates. (**L**) Schematic flowchart showing the Pten/PI3K/Akt/mTOR pathway and the different sets of double blocker experiments using LY294002/Torin1 in combination with SF1670. Scale bars represent 10μm in (**A, K**); the asterisk marks the injury site; GCL, ganglion cell layer; INL, inner nuclear layer; ONL, outer nuclear layer in (**A, K**); hpi, hours post injury; dpi, days post injury; Error bars represent SD; Ctl MO-Control morpholino (**A,B**); n.s., not significant.

**Figure EV6: Regulation of various genes by Mmp9 blockade and overexpression, effect of Notch signaling on Akt activation in zebrafish and on proliferation of MGPCs in zebrafish and mice retina, temporal and spatial expression profiles of *cfos* gene in injured retina.**

(**A**) The qPCR analyses of *adam10a*, *adam17a*, *rbpja* and *her4.1* in SB3CT-treated retina at 16hpi; *p < 0.01, n=6 biological replicates. (**B**) Densitometry plots for Western blots (corresponding to Figure 5J) show the regulation of Pten in DAPT-treated retinae at 2 and 4dpi; *p < 0.001; n=6 biological replicates. (**C** and **D**) Western Blot analyses of Akt, pAkt-Ser467, pAkt-Thr302 from retinal extracts prepared from retinae injected with different concentrations of DAPT at 4dpi (**C**), quantified by densitometry (**D**); *p < 0.001; n=6 biological replicates. Gapdh is the loading control. (**E**) Densitometry plots for Western blots (corresponding to Figure 5L) show the regulation of Pten upon *her4.1* knockdown in retinae at 2 and 4dpi; *p < 0.001; n=6 biological replicates. (**F** and **G**) IF microscopy images of retinal cross-sections show a concentration-dependent increase in the number of PCNA^+^ MGPCs in *her4.1* knockdown retina at 4dpi (**F**), which is quantified (**G**); *p < 0.02, n=6 biological replicates. (**H**) Densitometry plots for Western blots (corresponding to Figure 5M) show the regulation of PTEN, AKT, cMYC in DAPT-treated mice retinae at 2.5dpi; *p < 0.001; n=6 biological replicates. (**I** and **J**) IF microscopy images of retinal cross-sections show a concentration-dependent increase in the number of EdU^+^ MGPCs with DAPT treatment in mice retina compared to the control retina at 2.5dpi (**I**), which is quantified (**J**); Hoechst 3342 staining is the nuclear counterstain; *p < 0.0002; n=6 biological replicates. (**K**) Densitometry plots for Western blots (corresponding to Figure 5P) show the regulation of Pten in SB3CT-treated retinae at 16hpi and 2dpi; *p < 0.001; n=6 biological replicates. (**L**) The qPCR analyses of *ptena* and *ptenb* levels in *mmp9*-overexpressed retina at 2dpi; *p < 0.03, n=6 biological replicates. (**M**) Densitometry plots for Western blots (corresponding to Figure 5Q) show the regulation of Akt, pAkt-Ser467, pAkt-Thr302 in retinae treated with SB3CT alone and in combination with SF1670 at 4dpi; *p < 0.001; n=6 biological replicates. (**N**) Semi-quantitative PCR analysis of *cfos* gene expression at various time-points post-retinal injury. (**O**) BF microscopy images of retinal cross-sections show the mRNA *in-situ* hybridisation (ISH) of the *cfos* mRNA in the retina at 0.25, 0.5 and 1hpi. White arrowheads mark *cfos* mRNA signal. (**P**) BF and IF microscopy images of a retinal cross-section show the mRNA ISH of the *cfos* mRNA in the BrdU^+^ cells in the retina at 4dpi. White arrowheads mark co-labelled *cfos* mRNA and BrdU signal. Scale bars represent 10μm in (**F, I, O, P**); the asterisk marks the injury site and GCL, ganglion cell layer; INL, inner nuclear layer; ONL, outer nuclear layer in (**F, I, O, P**); hpi, hours post injury; dpi, days post injury. Error bars represent SD; Ctl MO-Control morpholino (**E-G**); n.s., not significant.

**Figure EV7: Schematic representation of role and regulation of Pten/PI3K/Akt/mTOR pathway and its interplay with Mmp9/Notch signalling during zebrafish retina regeneration.**

The model represents the regulation in MGPCs and the neighbouring cells of MGPCs.

**Table EV1. The list of primers used in this study.** First column describes primer name, second one is the ENSEMBL ID of the gene, and last one is the DNA sequence of the primer in 5’ to 3’ direction

